# Single-Cell Cross-Species Profiling identifies Conserved Transcriptional Networks in Early Pancreatic Tumourigenesis

**DOI:** 10.64898/2026.03.03.708839

**Authors:** Chiara Goossens, Colin lolos, Ana Lopez-Perez, Maurijn Kessels, Elisa Deom, Noella Bletard, Bernard Peers, Lydie Flasse, Marianne Voz

## Abstract

Pancreatic ductal adenocarcinoma (PDAC) is the most common form of pancreatic cancer and carries the poorest prognosis among all cancers, largely because it is frequently diagnosed at metastatic stages. It is therefore critical to identify reliable markers of preinvasive stages and to decipher the network driving preinvasive lesions to invasive carcinoma.

Here, we generated a zebrafish model in which KRAS^G12D^ is specifically expressed in pancreatic acinar cells, inducing acinar-to-ductal metaplasia that faithfully mirrors mammalian tumorigenesis. Single cell RNA-seq allowed us to capture transcriptional changes occurring at early stages of the disease. Cross-species comparison with mouse and human scRNAseq transcriptomes revealed a striking conservation of the genes upregulated during metaplasia, triggering common signalling pathways and regulatory programs. Notably, metaplastic cells reactivate a broad set of developmental genes expressed in multipotent pancreatic progenitors.

Mapping the acinar-to-cancer trajectories revealed a set of cytoskeletal and migration-related genes specifically upregulated during the late phase of metaplasia, immediately prior to malignant transformation, likely conferring invasive potential to these cells. SCENIC analysis further identified regulatory networks that become progressively activated as cells transition toward cancer, suggesting their involvement in the acquisition of malignant traits.

In conclusion, our cross-species comparison demonstrates a high degree of conservation in the molecular mechanisms driving pancreatic cancer progression from early to late stages across evolutionarily distant species, including zebrafish, mouse, and human, highlighting critical pathways that should be targeted to prevent cancer progression.

To allow researchers to easily explore gene expression profiles during pancreatic cancer progression across all three species, the datasets are publicly accessible via a user-friendly web platform (https://www.zddm.page.gd/)

## Background

Pancreatic ductal adenocarcinoma (PDAC) is the most prevalent form of pancreatic cancer and carries the worst prognosis of all common cancers. In 95% of cases, tumourigenesis initiation stems from the acquisition of an oncogenic mutation in the KRAS oncogene, followed by the accumulation of additional mutations, such as those in the tumour suppressor genes TP53, CDNK2A, SMAD4, and BRCA2 [1]. The acinar and the ductal cells are considered as the two main cell types of origin for this disease [2]. According to the current model, acinar cells undergo acinar to ductal metaplasia (ADM) and then progress into pancreatic cancer through Pancreatic Intraepithelial Neoplasia (PanIN) lesions. Ductal cells, on the other hand, are believed to progress to pancreatic cancer through Intraductal Papillary Mucinous Neoplasm (IPMN) lesions [3]. However, this model is overly simplified and needs further refinement. Indeed, murine ductal cells do not always advance through IPMN [4,5] and murine acinar cells do not consistently progress through preneoplastic PanIN stages to develop PDAC [6,7]. Furthermore, according to the model described by Marstrand et al., acinar cells do not directly transdifferentiate to ductal-like cells but rather first dedifferentiate into progenitor-like cells, which then differentiate into ductal-like cells [8]. This model is mainly based on the observation that during the acinar-to-ductal transition, several key factors involved in pancreatic embryonic development and progenitor specification—such as HNF1B, SOX9, PDX1, ONECUT1, SOX4, and GATA6—are upregulated. This capacity for dedifferentiation is also observed when human and murine acinar cells are transferred to *in vitro* culture, where they spontaneously acquire features characteristic of embryonic progenitors [9,10].

A detailed understanding of the initial stages of tumourigenesis is essential to identify early biomarkers capable of detecting tumours while still resectable. For this purpose, single-cell RNA sequencing (scRNA-seq) represents a method of choice. Indeed, by determining the transcriptomic profile of individual cells, it allows the identification of all transcriptional changes, tumour cells undergo along the paths leading to cancer. Schlesinger et al. pioneered this approach by applying scRNA-seq to model the transition from acinar to metaplastic cells, identifying over 3,000 genes differentially expressed during this early stage [11]. Identifying among all these transcriptional changes those that are truly driving the oncogenic process is a challenging task. Cross-species comparison of evolutionarily distant species such as humans and zebrafish provides an effective way to distinguish conserved, disease-relevant programs from species-specific features [12,13]. These comparisons have enabled the identification of “driver” mutations among all “passenger” mutations for several different cancers [14–16]. This cross-species approach was also very successful for melanomas where it highlighted the expected signatures relative to RAS and MAP kinase activation but also uncovered a new tumourigenic signature associated to neural crest cells formation[17]. High-throughput drug screening targeting this novel signature was performed in zebrafish and led to the identification of leflunomide as a new agent that inhibits melanoma cell growth [17–19]. These results highlight the power to combine cross-species comparisons to identify important drivers of the disease that are thereafter targeted by high throughput drug screening in zebrafish.

Currently, several zebrafish models of pancreatic cancer have been developed. In many cases, the *ptf1a* promoter is used to drive oncogenic Kras expression [20–23]. As a result, oncogenic Kras is activated at a very early developmental stage (around 34 hours post fertilisation (hpf)), in multipotent pancreatic progenitors before becoming restricted to the acinar lineage. Such activation at the embryonic stage does not reflect the human oncogenic process and prevents the precise identification of the tumour cell of origin. To address this limitation and generate a model with acinar-specific tumour origin, Park and Leach developed an inducible Cre-lox system to express *eGFP-KRAS^G12V^* in acinar cells at 3 weeks post-fertilization (*ptf1a*-CreERT2; *ubb*:LSL:GAL4-VP16; UAS:*eGFP-KRAS^G12V^*) [24]. This approach led to the development of lesions resembling human pancreatic intraepithelial neoplasia (PanIN) in only 2 of the 18 fish analysed. This low incidence of PanIN formation (∼11%), limits its utility as a robust pancreatic cancer model. Finally, Oh and Park also target the acinar lineage by driving *KRAS^G12D^* expression under the *ela3l* promoter (Ela3l:CRE; *ubb*:Lox-Cherry-Lox-GFP:*KRAS^G12D^*) [25]. Quite unexpectedly, the five tumours characterized in this study closely resembled pancreatic endocrine tumours, raising concerns about the model’s relevance for studying pancreatic ductal adenocarcinoma.

To establish an efficient acinar-derived model of pancreatic tumourigenesis, we used the *ela3l* promoter fused to GAL4 to induce a UAS:GFP-Kras^G12D^ transgene in mature acinar cells. Because *TP53* is mutated in more than 60% of human PDACs, we introduced these transgenes into a *tp53*-mutant background, which resulted in pancreatic tumour development in 100% of fish by one year of age. The first tumours emerged in three-month-old fish and exhibited ADM features that closely mirror those observed in mammalian models. We then performed single-cell RNA sequencing of zebrafish tumours and compared them with published mouse and human scRNA-seq datasets. This cross-species analysis revealed a remarkably high conservation of the transcriptional programs associated with metaplasia. By reconstructing the trajectory from acinar cells to cancer, we captured the continuous gene-expression changes occurring along this progression and identified a distinct set of genes specifically upregulated at the metaplasia–cancer boundary in the three vertebrate species. Finally, our SCENIC analysis uncovered several transcriptional programs that become progressively activated along the acinar-to-cancer trajectory, suggesting key roles in orchestrating the shift from metaplasia to malignancy.

## Methods

### Zebrafish maintenance, transgenic and mutant lines

Zebrafish (Danio *rerio*) were raised and cared according to standard protocols. The fish were maintained according to national guidelines and all experiments described were approved by the Ethics Committee of the University of Liège (protocol numbers 16-1851 and 21-2355). The tp53^zdf1^ mutant fish, referred to here as *tp53m*, carries a missense mutation (M214K) in the DNA-binding domain of p53, leading to loss of function [27]. This line was kindly provided by L. Zon and genotyped by sequencing following PCR amplification of genomic DNA. The Tg(ela3l:kaltA4) transgene was cloned into the pDESTol2CG2 vector using the Gateway cloning system; it contains 2979 bp of the ela3l promoter driving the expression of KalTA4, a fusion protein composed of the Gal4 DNA-binding domain and the TA4 activator domain [26]. The tg(UAS-e1b:GFP-Kras^G12D^) transgene was kindly provided by F. Argenton. For the control tg(UAS-e1b:GFP) transgene, the Kras gene was removed using the NEBuilder assembly method. All these transgenes have been then introduced into AB embryos by co-injection with the Tol2 transposase. Zebrafish were imaged using Leica microscope fluorescent binocular or on a Leica TCS Sp5 confocal

### Fixation, Sectioning, Stainings and Immunofluorescence

The tumours and control pancreata were fixed 24 hours with Neutral Buffered Formalin (NBF) 10% (HT501128, Sigma-Aldrich) at room temperature, washed with PBS, then conserved in ethanol 70% and embedded in paraffine by the GIGA-Immunohistology platform. Hematoxylin/eosin, eosin/alcian and Sirius red stainings were performed on 5 µm microtome sections by the GIGA immunohistology platform. Brightfield images were captured using the Zeiss Digital SlideScanner Axioscan 7. For the immunofluorescence, slides were deparaffinised and unmasked using EDTA buffer by the GIGA-Immunohistology platform. Endogenous peroxidase activity was blocked by 3% hydrogen peroxide (10 min at room temperature). Immunofluorescence staining was performed with anti-GFP (rabbit, Cell Signalling #2956S 1:200), anti-Caveolin-1(rabbit, Cell Signalling #3238 1:250), anti-α-Amylase (rabbit, Cell Signalling #3796 1:200), anti-Phospho-Histone H3 (pH3) antibodies (rabbit, Cell Signalling #3377T 1:200) overnight at 4°C followed by rabbit HRP SignalStain Boost (Cell Signalling #8114), and Tyramide-FITC, Tyramide-Cy3 (TSA Plus Cyanine 3 System #NEL744001KT Akoya BioSciences) or Tyramide-Cy5 substrate (TSA Plus Cyanine 5 System #NEL745001KT Akoya BioSciences). Slides were mounted in Prolong (Invitrogen) with DAPI 1:1000 and imaged using Zeiss Digital SlideScanner Axioscan 7. pH3 quantification was performed in QuPath by measuring the area occupied by pH3-positive cells relative to the DAPI-stained area in the pancreas [70].

### RNAscope experiments

Samples were fixed during 24 hrs at room temperature in 10% neutral-buffered formalin followed by standard paraffin embedding and sectioning. The RNAscope Multiplex Fluorescent v2 Assay (cat no. 323100, Advanced Cell Diagnostics, Newark, CA, USA) or the RNAscope 2.5 HD Assay-RED (cat no. 322350) was used according to manufacturer’s instructions with standard pretreatment conditions (15 min. Target Retrieval and 30 min. Protease Plus). Probes used were B. subtilis dapb (cat no. 320871) and Dr. Sox9b (cat no. 506291). Imaging was performed with an Axio Scan.Z1 (Zeiss, Jena, Germany) at 20x magnification. RNA in situ hybridization and imaging was performed by the VSTA core facility at VUB (https://vsta.research.vub.be). Statistical analysis was performed using the non-parametric Mann-Whitney t-test of the GraphPad Prism version 8.0.2 software.

### RNAseq of pancreatic multipotent progenitor cells

Pancreatic ventral bud cells were visualised using the transgenic line Tg(*ptf1a:GFP*) [71]. Four independent biological replicates with about 300 embryos at 35 hpf were micro-dissected and the tissue was mechanically dissociated. Cells were sorted based on their GFP expression on a FACS Aria II. cDNA synthesis was performed directly on the lysed cells using the SuperScript II reverse transcriptase (Invitrogen) and amplified with the KAPA HiFi HotStart Ready Mix (KAPA biosystems). Amplified cDNA was purified using Agencourt AMPure XP beads (Beckman Coulter, USA). The quality of the cDNA was verified by 2100 High Sensitivity DNA assay (Agilent Technologies) and the exact concentration of cDNA was determined using Quant-iT™ PicoGreen™ dsDNA Assay Kit (Invitrogen). 150 pg of cDNA were used as input to prepare the libraries using the Nextera XT DNA kit (Illumina). Sequences were obtained by the Genomic Platform at the GIGA (University of Liege) using the NextSeq500 Illumina Sequencer.

### ScRNAseq experiments

As soon as the tumours were visible, the fish were euthanized and the pancreatic tumours were dissected. To minimize potential sex-related heterogeneity, the scRNAseq experiments were performed exclusively on male fish. The dissected tissue was dissociated using a cocktail of enzymes (TrypLE Select 1x, 0,25 mg/mL Collagenase P, 0,35 mg/mL Collagenase IV) combined with a mechanical dissociation of 41 min at 37°C performed on the “gentleMACS™ Octo Dissociator with Heaters”. Dissociated cells were filtered with a 30 µm cell strainer (MACS SmartStrainers, Miltenyi Biotech), centrifuged 7 min at 300g and collected in HBSS^-^ 0.04% Ultrapure BSA. Then the viable cells labelled with calcein Violet 450 AM (Invitrogen, 15570597) were sorted by flow cytometry on BD FACSAria III to remove damaged cells and aggregated cells. For the control samples, 6 pancreata were dissected and pooled. The tissue was dissociated using TrypLE Select 1x, 0,04 mg/mL Proteinase K, 0,03 mg/mL Collagenase IV enzymes and a mechanical dissociation by pipetting up and down for about 20 min at 28°C, as healthy pancreas tissue digests much faster and the cells are more fragile compared to tumour tissue. Trypan blue was used to check viability on Countess 3 Cell Counter (Invitrogen) and then cells were diluted to about 1,2.10^6^ cells/mL. Single-cell RNA-seq libraries were generated using the 10x Genomics Chromium Single Cell 3′ v3 chemistry. Cells were loaded onto the Chromium Single Cell chip by the GIGA Transcriptomics platform, and the resulting libraries were sequenced on an Illumina NovaSeq 6000 instrument.

### RNA-seq data bioinformatical analyses

Raw reads were aligned to the zebrafish genome (GRCz11, Ensembl Release 103, ensembl.org) and quantified using the nf-core/rnaseq pipeline (v3.14.0) executed with Nextflow (v23.10.1) and the Docker profile. Normalization and differential expression analysis were performed using DESeq2 (v1.44.0) [72]; genes with absolute fold change ≥2 and adjusted p-value (padj) < 0.05 were considered differentially expressed.

### Single-cell RNA-seq data analyses

#### Read alignment and generation of gene–cell matrices

Sequencing output reads were provided as FASTQ files and aligned using the Cell Ranger pipeline (v6.1.2, 10x Genomics). Reads were mapped to their respective reference genomes supplemented with transgene sequences. For mouse samples, FASTQ files were obtained from the Gene Expression Omnibus (GEO) under accession number GSE141017 and aligned to the GRCm38 genome (Ensembl Release 102, ensembl.org) supplemented with the tdTomato sequence. For zebrafish samples, the GRCz11 genome (Ensembl Release 103, ensembl.org) was supplemented with GFP and Kras^G12D^ coding sequences. For the human dataset, raw count matrices and associated metadata were downloaded from the Genome Sequence Archive under project accession PRJCA001063.

#### Pre-processing, quality control, normalization and scaling

As we observed ambient RNA contamination in our zebrafish samples, we removed it by processing the raw gene–cell count matrices using the *remove-background* function from the CellBender package (v0.3.2) with default parameters. Doublets were identified and removed using the DoubletFinder R package (v2.0.6). All filtered count matrices (zebrafish, mouse, human) were imported into the Seurat R package (v4.4.0) for downstream analysis. Cells with fewer than 200 detected genes or with more than 10% mitochondrial gene expression were excluded. Genes expressed in fewer than three cells were removed from further analysis. UMI count matrices were log-normalized using Seurat’s NormalizeData function and scaled using ScaleData.

#### Data integration and batch correction

Batch correction was performed to ensure adequate overlap between biological conditions in the zebrafish dataset. Data integration was carried out using Harmony [32] with default parameters. In contrast, integration was not required for the mouse and human datasets, which were combined using a simple merge procedure.

#### Differentially expressed genes

We chose to use Seurat v4 rather than Seurat v5 to calculate our differentially expressed genes (DEGs) even though Seurat v4 tends to drastically underestimate the log2 fold change (log2FC), particularly for lowly expressed genes, because the ratio of cells expressing a gene in one population (pct1) versus the other (pct2) is not properly accounted for. In contrast, Seurat v5, while providing more accurate log2FC estimates, detects many more DEGs, including genes with minimal expression in metaplastic cells but absent in control acinar cells, which may lead to the detection of DEGs not truly relevant to the metaplastic process. This scenario is frequently encountered in our analyses, as control acinar cells express very few genes overall—the average number of unique transcripts per cell is 560, compared to 1,296 in metaplastic cells. This difference is largely due to the fact that over 50% of the acinar transcriptome is devoted to the expression of few acinar-specific enzymes [73]. Consequently, even when using Seurat v4, we identify a large number of upregulated genes and very few downregulated genes, aside from acinar-specific genes.

#### Identification of the Orthologous genes

To perform the cross-species comparison, zebrafish and murine differentially expressed genes (DEGs) were converted to their human orthologs using a self-compiled orthology table. Predicted orthologous relationships among zebrafish, mouse, and human were retrieved from Ensembl Release 103 and ZFIN. This table includes both one-to-one (1:1:1) orthologs as well as one-to-many relationships and is available upon request. Based on this orthology table, among the 2,724 upregulated genes identified in zebrafish, 2,372 genes (81%) had an ortholog detected in both the human and mouse datasets. Among the 4,948 upregulated mouse genes, 4,339 genes (87.7%) had an ortholog in both the human and zebrafish datasets. Finally, among the 4,958 upregulated human DEGs, 4,225 genes (85%) had orthologs detected in both zebrafish and mouse datasets. Only genes that had orthologs in the other species were considered for the Venn diagram analyses (Figure 5D) (i.e. 2372 zebrafish 4339 mouse and 4225 human genes). Among the 1,411 zebrafish genes comprising the developmental signature, 1,151 had an ortholog detected in both human and mouse datasets.

#### Gene ontology analyses

Gene set enrichment analysis (GSEA) [40] was performed with the list of all expressed genes ranked based on the fold change and the pct1/pct2 ratio between the metaplastic and acinar groups. Gene Ontology (GO) enrichment analyses were performed in R using the enrichGO function from the clusterProfiler package (v4.12.6). Differentially expressed genes were converted from gene symbols to ENTREZ IDs using species-specific annotation packages (org.Dr.eg.db for zebrafish, org.Mm.eg.db for mouse, and org.Hs.eg.db for human). Enriched GO terms with a q-value ≤ 0.05 (Benjamini–Hochberg correction) were considered significant and visualized using the enrichplot package.

#### Monocle and Scenic analyses

Trajectory analysis was performed with monocle 3 package and pseudotime was calculated for each cell [74]. Gene regulatory network analysis was performed using SCENIC [53]. Briefly, co-expression networks were first inferred using the GRNBoost algorithm to identify putative transcription factor–target gene modules. Motif enrichment analysis was then carried out with RcisTarget using the mc_v10_clust motif database and the motif2TF v10 annotation to identify significantly enriched motifs and retain direct TF–target interactions, thereby defining regulons. Finally, regulon activity in individual cells was quantified using AUCell.

#### Copy number variation estimation

Copy number variations (CNVs) were inferred from the mouse scRNAseq data using the inferCNV R package (v1.21.0, https://github.com/broadinstitute/inferCNV). Only tdTomato-positive cells (≥1 UMI) from the metaplastic clusters (prol-M, inter-M, late-M, and cancer) were analysed, with a gene expression cutoff of 0.1.

## Results

### The expression of GFP-Kras^G12D^ in acinar cells efficiently induces the formation of pancreatic tumours

To establish a zebrafish model of pancreatic cancer originating from acinar cells, we employed the GAL4/UAS system to drive the expression of KRAS^G12D^ specifically in acinar cells. Two transgenic lines have been generated (Figure 1A): a tg(ela3l:KalTA4; cmlc2:GFP) activator line where the *ela3l* promoter directs the expression of a KalTA4 fusion protein in acinar cells. The KalTA4 fusion protein contains the DNA binding domain of Gal4 fused to a TA4 activator domain [26]. The tg(UAS-e1b:GFP-KRAS^G12D^) effector line contains 14 UAS sequences upstream of a minimal *e1b* promoter driving the expression of a GFP-KRAS^G12D^ fusion protein [21]. In the double transgenic lines, referred to as Ac-K (for acinar KRAS^G12D^), the binding of the KalTA4 protein at the 14 UAS sites induces the expression of the GFP-KRAS^G12D^ protein in acinar cells. Fluorescence from GFP-KRAS^G12D^ was detected in these cells starting at 5 days post fertilisation (dpf) (Figure 1C). Of note, the fluorescence intensity observed with GFP-KRAS^G12D^ was noticeably reduced compared to that of unfused GFP (Ac-G), which was used as a control (Figure 1B). To confirm that GFP expression was restricted to the pancreatic acinar cells, we performed confocal imaging on whole larvae. In both models Ac-G and ac-K, GFP expression was restricted to the acinar cells (Figure 1D-E). These transgenes were introduced into the p53^M214K^ background, which carries a missense mutation in the DNA-binding domain of p53, resulting in its loss of function [27]. The fish were regularly monitored for the appearance of tumoral masses (Figure 1G) and euthanized immediately upon visual tumour detection. The dissected tumours were opaque and smooth (Figure 1I, yellow dashes) compared to healthy pancreata that appear more translucent and granular (Figure 1H, white dashes). In the p53^M214K/M214K^ (hereafter p53^m/m^) background, the first tumoral mass was detected at 110 dpf and 50% of the fish exhibit a visually detectable tumour by 197 days (Figure 1J). The p53 ^m/+^ fish develop detectable tumours at a slower rate with the first detectable tumour appearing at 138 dpf and 50% of the fish affected by 266 dpf. In absence of mutation for p53, fish also developed pancreatic tumours, but at a significantly slower rate (Figure 1J).

**Figure 1:**
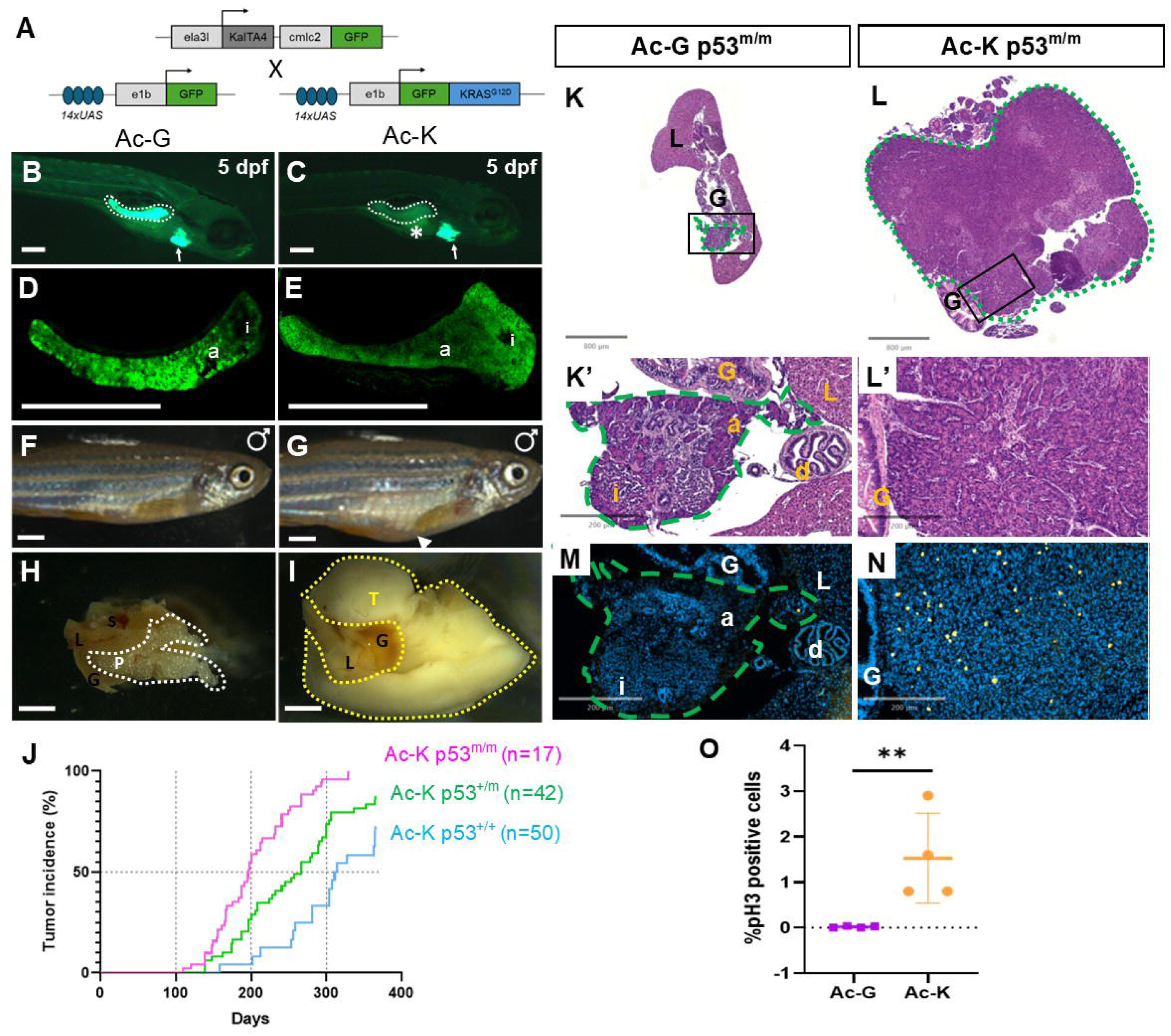
Generation and characterization of a zebrafish acinar-derived pancreatic tumour model. **(A)** Schematic representation of the transgenes used to generate the control (Ac-G) and the acinar-derived pancreatic tumour model (Ac-K). **(B-E)** 5-day zebrafish larvae expressing the protein GFP (**B,D**) or GFP-KRAS^G12D^ (**C,E**) in the acinar cells of the pancreas. White asterisk indicates gut autofluorescence. White arrows indicate the fluorescent heart, marked by the cmlc2:GFP reporter. Panels D and E show magnified views of the pancreata outlined with white dashed lines in panels B and C. i: principal islet, a: acinar cells. Scale bars : 200 µm. **(F-G)** Representative photographs of male zebrafish showing a marked abdominal protrusion, indicative of tumour presence (**G,** white arrow), compared to a control male (**F**). Scale bars: 2mm. **(H-I)** Dissected digestive systems of an adult control zebrafish (**H**), showing a healthy pancreas (P, white dashes) or of an Ac-K p53^m/m^ fish (**I**) with a pancreatic tumour (T, yellow dashes). L: liver, G: gut, S: spleen, P: pancreas, T: tumour. Scale bars: 2mm. **(J)** Tumour incidence across various *TP53* backgrounds. **K-L:** Hematoxylin and eosin (HE) staining of transversal sections of control pancreas (K) and Ac-K p53^m/m^ tumour (L) with close-up regions (K’,L’) outlined by black boxes. **M-N**: Immunofluorescence staining with phospho-histone H3 (pH3) antibody on control (M) and tumoral tissue sections (N). Control and tumoral pancreatic tissues are surrounded by green dashes. L; Liver, G; gut, d; extrapancreatic duct, i; principal islet, a; acinar cells. Scale bars: 200 µm. **O :** Quantification of pH3-positive cells in four Ac-K p53^m/m^ tumours compared to six control pancreas (3 p53^+/+^; 1 p53^+/m^, 2 p53^m/m^). The percentage of pH3 positive cells was calculated as the ratio of the pH3-positive area to the DAPI-positive area, based on the mean of two independent sections per fish. Each point represents an individual fish. Data are presented as mean ± SD; **P = 0.0095, determined using the Mann–Whitney test.

### Histological characterization of the pancreatic Ac-K p53^m/m^ tumours

Histological characterization of the Ac-K p53^m/m^ tumours using haematoxylin and eosin (HE) staining showed that tumours occupy a large portion of the peritoneal cavity in the fish in contrast to the healthy pancreas in control (Figure 1K-L). To assess the proliferation rate of tumour cells, we performed immunofluorescence staining using a phospho-histone H3 (pH3) antibody which specifically marks nuclei in M phase. While very few mitotic cells are detected in the control pancreas (0.03%), 1.5% of the tumour cells were in mitosis, indicating a 50-fold increase in proliferation rate (Figure 1 M-O).

Further examination of HE-stained sections revealed that the tumours exhibit disrupted tissue architecture compared to control acinar cells. In healthy tissue, acinar cells are arranged around a central lumen to form an acinus (outlined with white dashed lines in Figure 2A). They display a characteristic pyramidal shape with a large, round nucleus located at the basal pole (Figure 2A,B). In contrast, tumoral cells fail to form organized acini (Figure 2C) and lack their typical pyramidal morphology (Figure 2D). Moreover, whereas normal acinar cells contain abundant apically located zymogen granules (Figure 2B), tumoral cells exhibit a marked reduction in visible zymogen granules (Figure 2D). These findings suggest a reduced expression of digestive enzymes in tumour cells.

**Figure 2:**
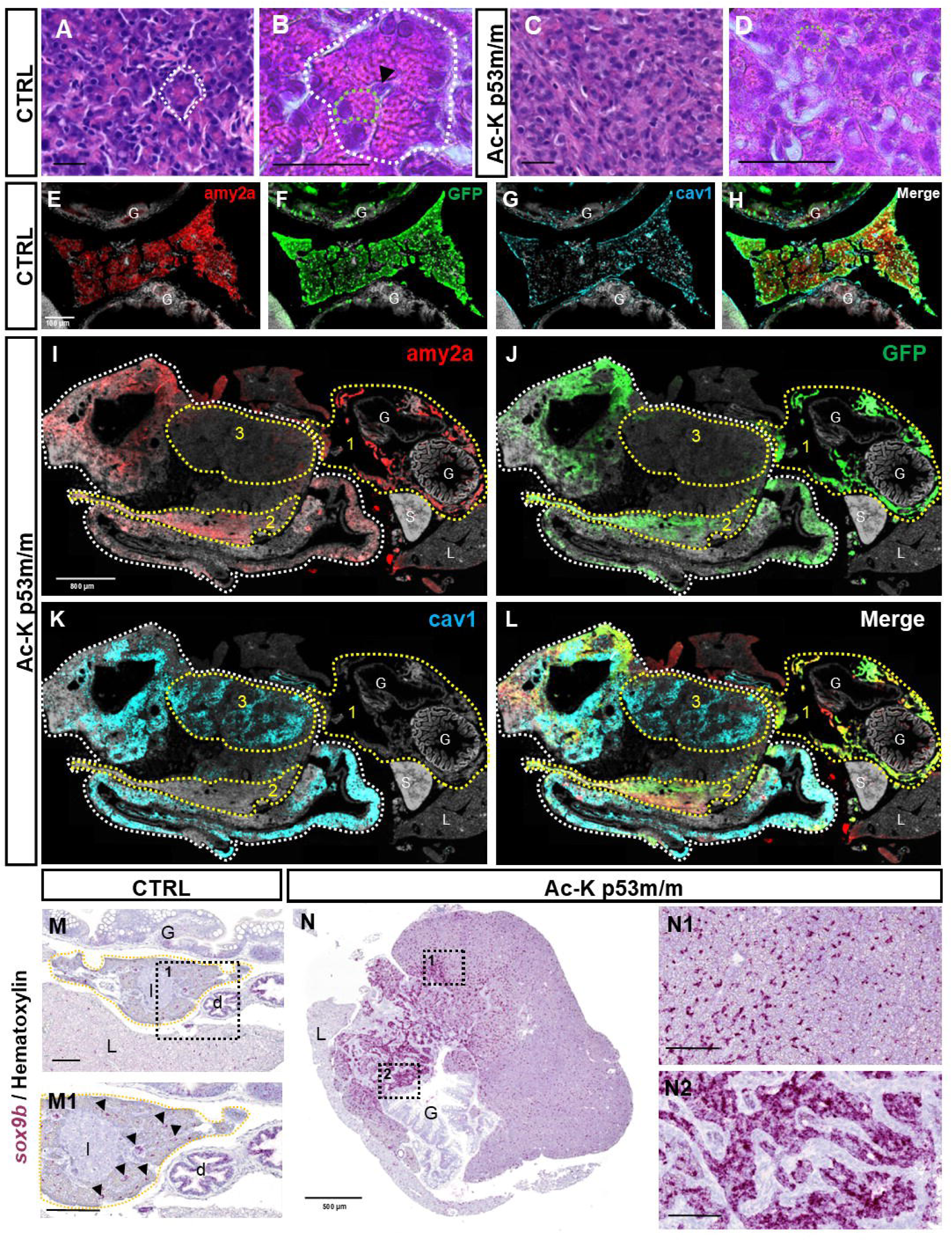
Ac-K p53m/m tumours display disrupted acinar architecture and metaplastic features. **A–D**: Hematoxylin and eosin (HE) staining of healthy control pancreas (A–B) and Ac-K p53m/m tumour tissue (C–D) showing that acinar cell architecture and morphology are disrupted Ac-K p53m/m tumour. (A) Control pancreatic tissue is primarily composed of acinar cells organized around a lumen to form acini (an example outlined with white dashed lines). (B) Acinar cells contain a round nucleus located at the basal pole and an eosinophilic cytoplasm, reflecting the high concentration of zymogen granules at their apical pole. A representative acinar cell is outlined with green dashed lines. Centroacinar cells are visible within the acinar lumen (black arrowhead). (C) Tumour tissue is composed of disorganized cells. (D) Tumour cells display reduced zymogen granules and loss of cell polarization. These observations were consistent across 15 Ac-K p53^m/m^ tumours and 5 healthy control pancreata (3 p53+/+, 1 p53+/m, 1 p53m/m).) G: gut; L: liver; d: ducts; I: islet. Scale bars = 20 µm. **E-L:** Immunofluorescence staining of control (E-H) and Ac-K p53^m/m^ tumours (I-L) for Amy2a (in red), GFP (in green) and Cav1 (in turquese) and DAPI in grey. Amy2a expression is reduced in tumoral tissue (I), concomitantly with GFP expression (J), while Cav1 expression was increased (K). These results were obtained in 2 metaplastic p53^m/m^ tumours and 5 controls (1 p53^+/+,^; 2 p53^+/m^, 2 p53^m/m^). **M-N:** *sox9b* mRNA detection by RNAScope on healthy (M) and Ac-K p53^m/m^ tumoral pancreas (N). (M) *sox9b* staining is detected in ducts (black arrows, M1) while in tumours sox9b is also detected in the tumoral tissue in isolated cells (N1) or in nearly all cells (N2). These results were obtained in 2 metaplastic p53^m/m^ tumours and 2 p53^+/+,^ controls. Scale bars =100 µm except for I-L (800µm) and N (500µm). Control pancreatic exocrine pancreas tissue is outlined by yellow dashes while tumoral tissue by blue dashes. G: gut, L: liver, d : duct, I : islet.

To explore this further, we performed immunohistochemistry for the acinar enzyme amylase (Amy2a). In control tissue, Amy2a expression was strong in all acinar cells (Figure 2E). In contrast, tumoral samples showed heterogeneous expression, with regional variation in staining intensity (Figure 2I). For example, tumour 21HIS2007 displayed a “healthy” zone with preserved histological architecture that surrounds the digestive tract, exhibiting high Amy2a expression comparable to control tissue (zone 1 in Figure 2I). However, adjacent tumoral regions (outlined by a white dashed line) showed area with reduced Amy2a expression (e.g., zone 2) as well as a larger region exhibiting an almost complete loss of Amy2a signal (e.g., zone 3). A similar pattern was observed for GFP (Figure 2J), which showed a concomitant decrease with *amy2a* in the tumoral region (zone1 > zone 2 > zone 3). This reduction in GFP is likely due to downregulation of *ela3l* acinar promoter activity in tumoral cells, leading to decreased KalTA4 expression and, consequently, reduced GFP-Kras^G12D^ expression. This decrease in GFP was also evident during tumour dissections, where 36% of the tumours (57 out of 158) exhibited complete or near-complete loss of GFP fluorescence (Supplementary Figure 1). Notably, reduced expression of Amy2a and GFP was frequently accompanied by a marked upregulation of Caveolin1 (Cav1) (figure 2K), which has been reported as a specific marker of acinar-derived tumours in mice [4]. Cav1 expression was particularly high in zone 3, where Amy2a and GFP expression were markedly reduced. These data indicate that Cav1 increases as Amy2a and GFP expression declines, suggesting that Cav1 may serve as a useful readout of metaplasia progression. We next tested whether the loss of acinar features was accompanied by an increased expression of ductal markers such as *sox9b*, a key metaplastic player in mice [28,29]. RNAscope experiments revealed that in control pancreas, *sox9b* expression is restricted to pancreatic ducts (black arrowheads in Figure 2M1) while ectopic *sox9b* expression is observed in the tumoral tissue (Figure 2N). Tumour heterogeneity is also observed here, with regions where *sox9b* is expressed in isolated cells (Figure 2N1), while in other areas, nearly all cells express *sox9b* (Figure 2N2). Similar results were obtained using fluorescent RNAScope (Supplementary Figure 2).

In conclusion, our data strongly suggest that expression of KRAS^G12D^ in acinar cell induce a metaplastic process, characterized by a loss of acinar traits and the emergence of ductal features.

### A Subset of Tumours Progresses to Invasive PDAC

Among the 17 tumours characterized by HE staining and immunohistochemistry, the majority (15 out of 17) exhibited a similar metaplastic histological profile (Table 1A, col. 3). However, two tumours progressed to pancreatic ductal adenocarcinoma (PDAC), exhibiting exhibiting invasion of surrounding tissues as well as local tissue infiltration of the pancreatic parenchyma. Specifically, tumour 21HIS478 invaded the gut (Figure 3A, A’) while tumour 21HIS474 invaded both the gut (Figure 3B, green dashed outlines) and liver (Figure 3B’). To characterize more tumours at advanced stages, we selected 14 additional samples that showed a complete or near-complete loss of GFP fluorescence at the time of dissection. Among these, eight tumours displayed PDAC characteristics such as nuclear atypia and varying degrees of glandular differentiation (Figure 3 C-E). While three of these PDAC exhibited invasion restricted to the pancreatic parenchyma, five also invaded neighbouring organs (Table 1B, col. 5). All 10 PDAC were classified by expert clinical pathologists according to the human WHO histological classification. Two tumours exhibited well-defined duct-like glandular structures, typical of well-differentiated PDAC (Figure 3C). Four others exhibited less well-defined glands with formation of cribriform structures and were classified as moderately differentiated (Figure 3D). The remaining four showed poorly formed glands and were classified as poorly differentiated (Figure 3E) (Table 1, col.3). As mucin production further supports PDAC classification in humans [30], we performed Alcian Blue (AB) staining to assess mucin expression. Strong AB staining was detected in the ductal-like structures of well-differentiated PDAC as reported for this histological grade (Figure 3F). Moderately differentiated tumours showed mild AB staining (Figure 3G), while poorly differentiated tumours displayed no detectable AB staining (Figure 3H), also in line with [30] (Table 1, col. 6).

**Figure 3:**
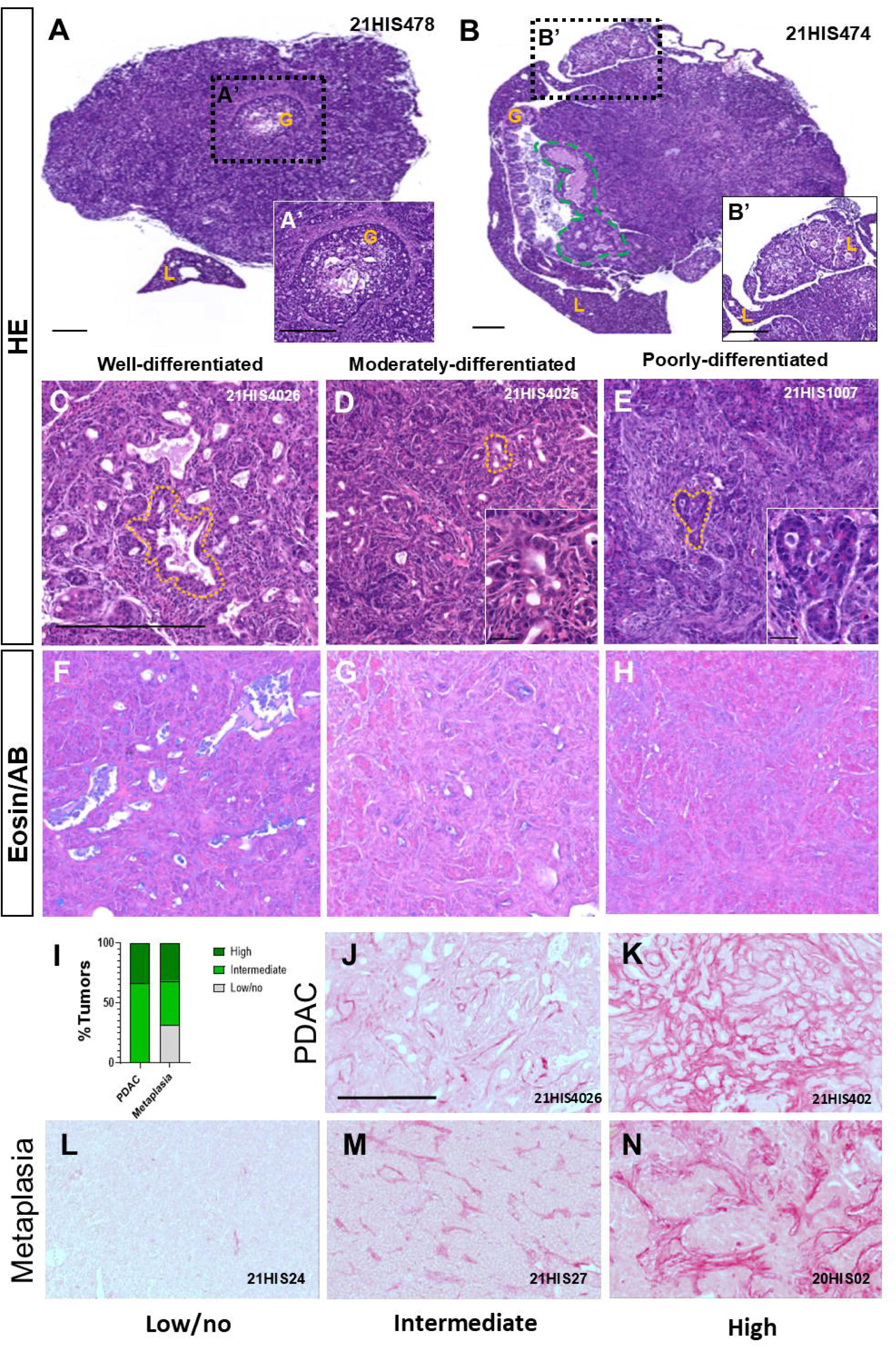
Histomorphological PDAC classification and associated desmoplasia across PDAC and metaplastic stages. **A-B:** H&E staining of moderately differentiated PDAC showing invasion into the gut (A, A’) or into both the gut (B, green dashed outlines) and liver (B, B’). **C-H:** HE and eosin alcian blue (Eosin/AB) stainings of well-differentiated (C, F), moderately differentiated (D, G) and poorly differentiated (E, H) PDAC. One gland is highlighted in C-D with yellow dashed lines and is magnified in the inset. L: liver, G: gut. **I.** Qualitative assessment of Sirius Red staining in PDAC and metaplastic tumours, categorized as high, intermediate, or absent. **J–N.** Representative Sirius Red stainings of PDAC (**J–K**) and metaplastic tumours (**L–N**). All PDAC samples displayed desmoplasia, with either intermediate (**B**, *n* = 6) or high (**C**, *n* = 3) staining levels. Among metaplastic tumours, Sirius Red staining was absent in some cases (**D**, *n* = 6), while others exhibited intermediate (**E**, *n* = 7) or high (**F**, *n* = 6) collagen deposition. The sample identification number of the tumor is indicated on each section. Scale bars: 250uµm (A–H), 200uµm (J–K) J

**Table 1:** Overview of the histological characteristics of Ac-K p53^m/m^ tumours. Tumours were either randomly selected (A) or selected based on reduced GFP levels (B). Column 1 indicates the sample identification number and whether the sample was used for scRNA-seq experiments (PT1-PT6). Column 2 reports the age of the fish (months and days) at euthanasia due to tumour presence. Column 3 describes tumour histology after H&E staining: Meta, metaplasia; well-diff. PDAC, well-differentiated PDAC; mod.-diff. PDAC, moderately differentiated PDAC; poor-diff. PDAC, poorly differentiated PDAC. Column 4 indicates secondary histology when tumours displayed two distinct H&E-stained regions. Column 5 specifies sites of invasion (liver or gut), local invasion of the pancreatic parenchyma, or absence of detectable invasion. Column 6 indicates the presence or absence of Alcian blue staining. Column 7 reports Sirius Red staining as absent (−), intermediate (+), or high (++); an asterisk indicates tumours exhibiting focal rather than widespread staining. ND, not determined.

|  | 1 | 2 | 3 | 4 | 5 | 6 | 7 |
| --- | --- | --- | --- | --- | --- | --- | --- |
|  | ID | Age | Histology | Secondary histology | Invasion | Alcian blue staining | Sirius red staining |
| A) Randomly selected tumours |  |  |  |  |  |  |  |
| 1 | 18HIS3784 | 4m16d | Meta | / | No | No | + |
| 2 | 19HIS1862 | 4m16d | Meta | Healthy | No | No | ++ |
| 3 | 21HIS1006 | 4m20d | Meta | Healthy | No | No | + |
| 4 | 19HIS3627 | 4m26d | Meta | / | No | No | ND |
| 5 | 19HIS3630 | 6m13d | Meta | / | No | No | ++ * |
| 6 | 20HIS0200 (PT1) | 6m18d | Meta | / | No | No | ++ |
| 7 | 20HIS0203 (PT2) | 8m3d | Meta | / | No | No | + |
| 8 | 21HIS467 | 8m15d | Meta | Healthy | No | No | ++ * |
| 9 | 21HIS468 | 8m17d | Meta | Healthy | No | No | - |
| 10 | 21HIS465 (PT3) | 8m17d | Meta | / | No | No | - |
| 11 | 21HIS471 | 8m23d | Meta | / | No | No | - |
| 12 | 21HIS472 | 8m23d | Meta | / | No | ND | ++ |
| 13 | 21HIS474 | 9m6d | Mod. diff. PDAC | / | Gut and liver | Yes | ++ |
| 14 | 21HIS475 | 9m6d | Meta | Healthy | No | ND | - |
| 15 | 21HIS476 | 9m13d | Meta | Healthy | No | ND | - |
| 16 | 21HIS477 | 9m13d | Meta | / | No | ND | ++ |
| 17 | 21HIS478 | 9m24d | Mod. diff. PDAC | Meta | Gut | Yes | ND |
| B) GFP reduced tumours |  |  |  |  |  |  |  |
| 1 | 21HIS1007 | 4m29d | Poor. diff. PDAC | Meta | Liver | Yes | ++ |
| 2 | 21HIS2007 | 5m22d | Meta | / | No | No | - |
| 3 | 21HIS1014 | 6m12d | Well diff. PDAC | Meta & Healthy | Pancreatic parenchyme | Yes | + |
| 4 | 21HIS2700 | 7m13d | Poor. diff. PDAC | Meta | Gut | ND | + |
| 5 | 21HIS2701 | 7m13d | Mod. diff. PDAC | Meta | Gut | Yes | + |
| 6 | 22HIS2482 (PT4) | 7m29d | Meta | / | No | ND | + |
| 7 | 21HIS4025 | 8m2d | Mod. Diff. PDAC | Meta | Pancreatic parenchyme | Yes | ++ |
| 8 | 21HIS4026 | 8m2d | Well diff. PDAC | Meta | Pancreatic parenchyme | Yes | + |
| 9 | 22HIS2484 (PT5) | 8m5d | Meta | / | No | ND | + |
| 10 | 21HIS2011 | 8m7d | Poor. diff. PDAC | Meta | Liver | No | + |
| 11 | 22HIS2837 (PT6) | 8m20d | Meta | Healthy | No | ND | + |
| 12 | 21HIS2702 | 8m24d | Meta | / | No | ND | + |
| 13 | 21HIS2703 | 8m24d | Meta | / | No | ND | ND |
| 14 | 21HIS2704 | 9m16d | Poor. diff. PDAC | / | Gut | Yes | + |

Given that desmoplasia—a fibrotic response marked by abundant extracellular matrix (ECM) and dense collagen-rich stroma—is a hallmark of PDAC [31], we examined its presence in our zebrafish PDAC tumour samples. To this end, we performed Sirius Red staining, which specifically highlights collagen fibres. All nine PDAC samples analysed exhibited desmoplasia, albeit with varying intensities (Figure 3 I-K, Supplementary Figure 3). These differences in desmoplastic response did not correlate with the PDAC histological classification (Table 1, col.3 & col.7). Desmoplasia was also observed in metaplastic tumours, with six samples (out of 19) showing high levels of fibrosis (Figure 3N, Supplementary Figure 4A-F). An additional seven tumours exhibited intermediate levels of Sirius Red staining (Figure 3 I,M; Supplementary Figure 4, G–M), whereas no detectable staining was observed in the six remaining samples (Figure 3 I,L; Supplementary Figure 4, N–S). These findings, summarized in Table1 (col. 7), indicate that while desmoplasia is a consistent feature of PDAC in zebrafish, it can also arise during metaplastic stages.

### Molecular Characterization of Pancreatic Tumourigenesis through scRNAseq Analysis

To molecularly characterize the different steps of pancreatic tumourigenesis, we performed scRNA-seq on six tumours resected at different time points (6 to 9 months) and on two independent pooled control samples, each generated from six healthy pancreata (see Methods). For each tumour sample, one portion was processed for scRNA-seq and the other for histological analysis. The histology indicated that the tumours were composed mainly of metaplastic cells, with no evidence of more advanced stages (Table 1). In total, 34186 cells from the six tumours and 9281 cells from the healthy pancreata passed all quality control criteria (see “MM”, Supplementary Figure 5). These cells were integrated using Harmony [32] and clustering resulted in the identification of 11 clusters (Figure 4A). Because zebrafish cell-type signatures remain poorly defined, we assembled a reference table of specific markers from the literature and from single-cell developmental atlases [33,34] (Supplementary Figure 6). Using this marker set, we assigned cell identities to all clusters. DotPlot analysis using a representative subset of markers confirmed the accuracy of these assignments. (Figure 4F). Four clusters correspond to cells of the microenvironment such as fibroblast (cl6), macrophages (cl5), T-lymphocytes (cl7) and neutrophils (cl9). Cluster 8 was identified as pancreatic ductal cells and the cluster 10 contain both endocrine and tuft-like cells.

**Figure 4:**
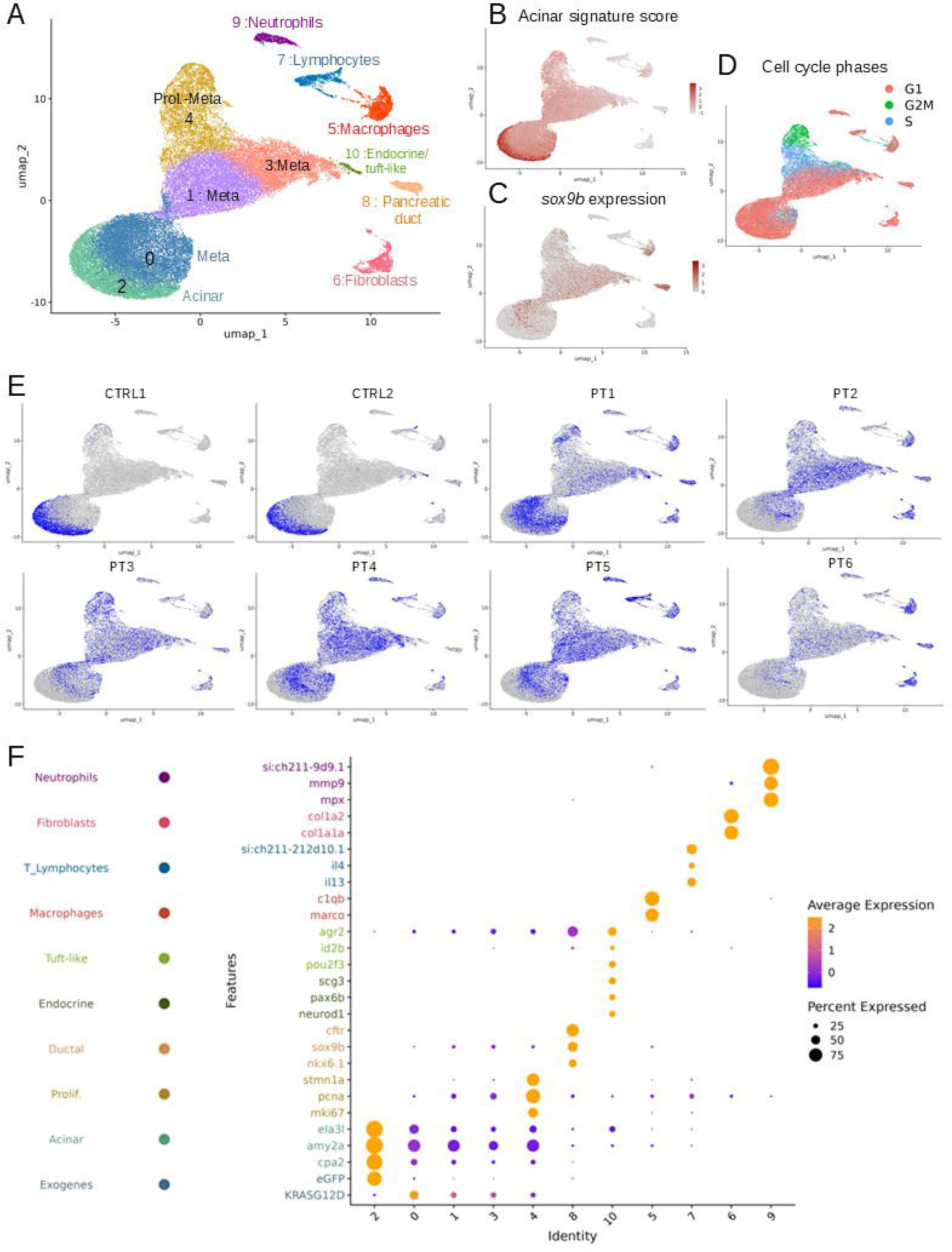
Transcriptomic profiling of Zebrafish Pancreatic Tumours by scRNA-seq. Cells were obtained from six tumour samples (PT1–PT6) and two healthy pancreas samples as controls (CTRL1–2) **A.** UMAP representation of all cells grouped into distinct clusters. **B-D.** UMAP plots showing the acinar signature score (B), *sox9b* expression levels (C) and cell-cycle status, with cells in G1 (red), S (blue) and G2/M (green) (D). **E.** UMAP plots showing the contribution of each sample, with cells from the highlighted sample in blue and all other cells in grey. **F.** Dot plot showing the expression of cell type–specific markers across clusters. The dot size indicates the proportion of cells expressing the gene, while the color intensity reflects the average expression level among those cells.

Cluster 2 represents healthy acinar cells, characterized by a high acinar signature score (Figure 4B) driven by elevated expression of digestive enzymes such as *ela3l*, *cpa2*, and *amy2a* (Figure 4F). This cluster is predominantly composed of cells from the two healthy control samples (Figure 4E; Supplementary Figure 7A). Four clusters (cl 0, 1, 3, and 4) likely correspond to tumoral metaplastic cells, as indicated by a marked downregulation of the acinar gene signature (Figure 4B,F) together with an upregulation of the ductal and metaplastic marker *sox9b* (Figure 4C,F). Additionally, we observed that metaplastic cluster 4 exhibit proliferative features as indicated by the expression of *mki67, pcna* and *stmn1a* (Figure 4F). The basal region of cluster 4 exhibits an S-phase gene expression signature, whereas the upper region is enriched in G2/M-phase markers (Figure 4D). This cluster represents 17,6% of all metaplastic cells.

To more precisely capture the transcriptional changes associated with metaplasia, we refined our analysis by subclustering pancreatic exocrine cells, thereby excluding all microenvironmental cell types. This approach revealed five distinct clusters of metaplastic cells, including two proliferative clusters corresponding to S-phase and G2/M-phase signatures (Supplementary Figure 8A). Metaplasia appeared to progress from acinar cells toward clusters 0, 1, and 3, as evidenced by the gradual loss of acinar markers and the concomitant increase in the metaplastic markers *krt94* (the ortholog of *KRT19*) and *cav1* (Supplementary Figure 8B). Cav1 was visualized by IHC as a marker of metaplasia progression (Figure 2K). Differential expression analysis was conducted in Seurat v4, comparing the pool of cells from intermediate and late metaplastic clusters with healthy acinar cells. Early metaplastic cells were excluded to avoid attenuating the magnitude of differential expression by including cells with weaker transcriptional changes. The proliferative clusters were also omitted to ensure that the gene list reflected metaplastic programs rather than proliferation-associated transcripts. Indeed, differential expression analysis comparing the two proliferative with the three non-proliferative metaplastic clusters revealed enrichment not only for classical S-phase and G2/M-phase gene signatures but also for multiple proliferation-associated genes, including histones and other cell cycle regulators. Our differential expression analysis identified 2800 differentially expressed genes (DEG) (absolute log₂ fold change (FC)> 0.25; minPCT>0,1, FDR < 0.001), with 2724 genes upregulated and 76 downregulated (Figure 5A and Additional file 1, sheet “dr_DEG”). Among these downregulated genes, acinar digestive enzymes are particularly represented, with 10 enzymes showing drastic reduction. Notably, *ela3l* emerged as the most downregulated gene (Log2FC = -7.3) likely explaining the loss of fluorescence observed in many tumours, since GFP–Kras expression is driven by the *ela3l* promoter. Consistent with the IHC shown above (Figure 2I), *amy2a* is also drastically downregulated in the metaplastic cells (Log2FC of -4.0). Among the positively regulated genes, *olfm4.1* and *olfm4.2* are the two top genes, with a log2FC of 6.4 and 6.8, respectively. OLFM4 has been shown to be a key mediator of STAT3 signalling in Human Hepatocellular Carcinoma [35], a pathway critically involved in the metaplastic process [36]. Also, among the top up-regulated genes, we found a huge number of genes involved in translation such ribosomal proteins *rps* and *rpl* and several eukaryotic translation initiation factors genes (Additional file 1, sheet “dr_DEG). This enrichment is likely due to the increased demand for protein synthesis to sustain uncontrolled cell growth in cancer cells [37].

**Figure 5:**
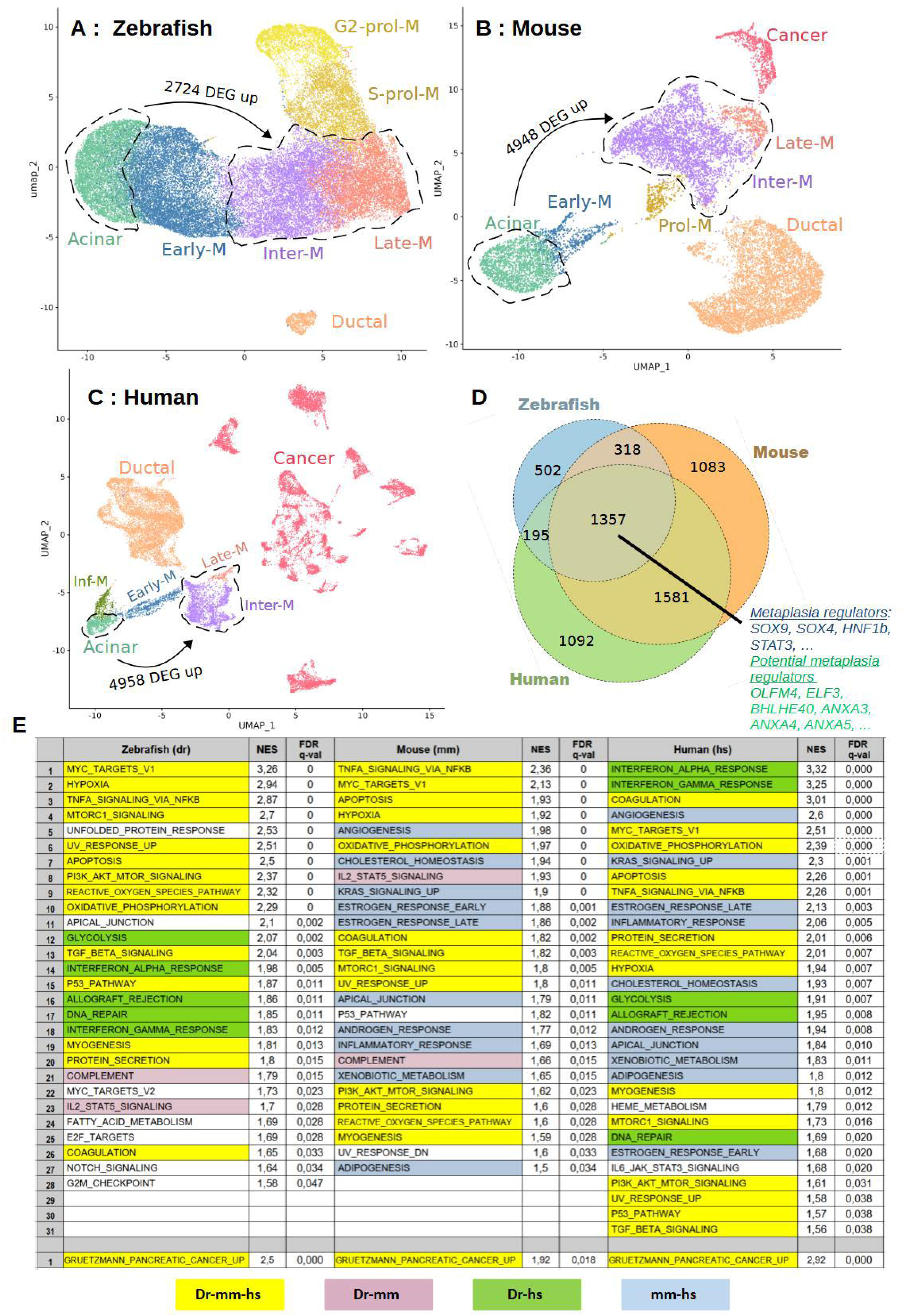
Cross-species comparison of the transcriptome of metaplastic cells. “A-C: UMAP plots of zebrafish (A), murine (B), and human (C) scRNA-seq data, grouped based on their characteristics as defined in Supplementary Figures 9 and 10 **D**. Venn diagram showing upregulated DEGs in metaplastic (intermediate and late) versus acinar cells across zebrafish, mouse and human species. **E**. GSEA analysis of pathways significantly enriched in zebrafish, murine and human metaplastic cells. NES : normalized enrichment score

### Cross-Species Analysis Reveals Highly Conserved Pathways in the Metaplastic Process

To uncover transcriptional changes critical to disease progression from this large set of upregulated genes, we performed a cross-species comparative analysis of metaplastic cells, combining zebrafish, murine, and human datasets and focusing on genes consistently upregulated across all three species.

For the murine data, we selected the study of Schlesinger et al., who analysed by scRNAseq the progression of tumoral cells from preinvasive lesions to cancer [11]. In this study, Kras^G12D^ was induced in acinar cells by tamoxifen administration to six- to eight-week-old Ptf1a-Cre^ER^, LSL-Kras^G12D^, LSL-tdTomato mice. Pancreata were collected for single-cell isolation at six time points after tamoxifen induction (17 days, 6 weeks, 3, 5, 9 and 15 months). We downloaded the raw data and analysed it using a similar pipeline as for zebrafish to enable an optimal comparison across the three species (see Methods). After exclusion of microenvironmental cells, pancreatic cells were grouped into 15 clusters as shown in the UMAP plot (Supplementary Figure 9A). These include acinar cells (cluster 0), ductal cells (clusters 1, 2, 4, and 12) and tuft cells (cluster 14) (Supplementary Figure 9A-B). Eight clusters (3, 5, 7–11, and 13) are classified as metaplastic cells, as they show a marked reduction in acinar marker expression alongside induction of the ductal markers *Krt19* and *Sox9*, as well as the reported metaplastic markers *Onecut2*, *Foxq1*, and *Runx1* (Supplementary Figure 9C). Within this metaplastic compartment, cluster 10 exhibits proliferative properties, while cluster 7 represents early metaplastic cells, as it begins to express the ductal markers *Krt19* and *Sox9* while remaining *Cpa1* and *Cel* positive (Supplementary Figure 9C). In contrast, cluster 6, derived nearly exclusively from 15-month-post injection mouse that developed invasive PDAC, corresponds to the cancer state, as it exhibits high levels of copy number variations (CNVs), (Supplementary Figure 9D), as reported [38]. Cluster 8—also originating from 15-month post injection mouse – show very few CNV aberrations (Supplementary Figure 9D) and was therefore proposed to represent the pre-cancerous tissue surrounding PDAC lesions [38]. We therefore classified cluster 8 as late metaplastic. The annotation of all clusters is shown in Supplementary Figure 9B.

Differential expression analysis comparing intermediate and late metaplastic cells to healthy acinar cells, excluding proliferative and early metaplastic clusters (Figure 5B), as done for the zebrafish dataset identified 4948 upregulated and 76 downregulated genes (absolute log2FC > 0.25, min.pct > 0.1, FDR < 0.001; Additional file 1, sheet “mm_DEG”).

For the human dataset, we selected the study by Peng et al., who performed scRNA-seq on PDAC samples from 24 patients and on pancreatic tissue from 11 control individuals [39]. We analysed the human data using a pipeline similar to that used for the zebrafish and mouse datasets (see Methods). After exclusion of microenvironmental cells, pancreatic cells, were grouped into 20 clusters as shown in the UMAP plot (Supplementary Figure 10A). Cells derived from healthy control pancreas samples are mainly distributed across four distinct clusters: two ductal clusters (clusters 0 and 13) and two acinar clusters (clusters 8 and 15) (Supplementary Figure 10C). Cluster 15 express acinar markers but also initiate ductal marker expression such as SOX9 and KRT19 (Supplementary Figure 10F), suggesting that these control-derived cells may have undergone a metaplasia-like process. Such non-oncogenic metaplasia is well documented and commonly observed in the context of pancreatic inflammation [8]. On the other hand, PDAC-derived cells were distributed across two major regions that we annotated as “cancer” and “pre-cancer” clusters (Supplementary Figure 10D). Indeed, the 13 “cancer” clusters were previously grouped into a single cluster called “*ductal cell type 2”* (Supplementary Figure 10E) which was reported to exhibit extensive CNV alterations indicative of an advanced cancer state [39]. In contrast, the “pre-cancer” clusters (clusters 9 and 1) were annotated as “ductal cell type 1” and “acinar (Suppl. Figure 10D-E). These clusters were shown to exhibit very low levels of CNV aberrations [39], suggesting that they represent precancerous tissue in the vicinity of PDAC lesions. Marker analysis indicates that these “pre-cancer” clusters consist of metaplastic cells. Indeed, these cells show i) reduced expression of acinar markers, ii) increased expression of the ductal genes *KRT19* and *SOX9* iii) increased expression of key actors of metaplasia such as *SOX4* and *HNF1B* iv) increased of metaplastic markers previously identified in mice such as *ONECUT2, FOXQ1*, and *RUNX1* (Supplementary Figure10F). Among these, cluster 9 was classified as early metaplastic, as it retained partial expression of acinar markers and exhibited a modest induction of ductal and metaplastic markers (Supplementary Figure10F). In contrast, cluster 14a was classified as late metaplastic, displaying a strong reduction in acinar marker expression together with the highest levels of metaplastic markers (Supplementary Figure 10F). Importantly, this cluster seems to represent a transitional state between metaplastic and cancerous cells, as the second subcomponent of this cluster (cluster 14b) is positioned centrally among all other cancer cell populations (Supplementary Figure 10A).

Differential expression analysis comparing intermediate and late metaplastic cells to healthy acinar cells (Figure 5C) identified 4,958 upregulated and 119 downregulated genes (absolute log2FC > 0.25, min.pct > 0.1, FDR < 0.001; Additional file 1, sheet “hs_DEG”). As in zebrafish and mouse, the downregulated genes were predominantly acinar digestive enzymes.

To identify metaplastic genes common in the three vertebrate species, we converted the zebrafish and murine DEG genes into their human orthologs (see Methods). A cross-species comparison of upregulated genes revealed a remarkably high level of conservation, with 1374 genes shared across all three species (Figure 5D, Additional file 1, sheet “common_DEG_UP”). This overlap was significantly greater than expected by chance, as assessed by a hypergeometric test (p < 10⁻¹). Among the common genes were well-established regulators of metaplasia including *SOX4*, *SOX9*, and *HNF1B.* Interestingly, our cross-species comparison also highlights a broad set of genes, including *SOX11,* ELF3*, ELF1, ETS2* and *BHLHE40,* whose roles in metaplasia have not yet been described and therefore warrant further investigation. Furthermore, cross-species comparison of downregulated genes reveals that the consistently repressed genes encode pancreatic enzymes. Together, these results reveal a conserved process across all three species: a reduction in pancreatic enzyme expression accompanied by upregulation of metaplastic genes

To identify pathways critical for the metaplastic process, we performed gene set enrichment analyses (GSEA) [40], initially focusing on the 50 hallmark gene sets, that represent core biological processes and signalling pathways in a non-redundant manner. Remarkably, most pathways significantly enriched in zebrafish (FDR<0,05) were also significantly enriched in mouse and human (highlighted in yellow in Figure 5E). Among these, three pathways are known to be activated by constitutively active KRAS, the PI3K/Akt/mTOR, the Raf/MEK/ERK/Myc and the NF-κB pathways, whose activation favours oncogenic transformation [41]. We also identified the TGF-β pathway, shown to induce acinar-to-ductal metaplasia and to establish a permissive context for the emergence of oncogene-driven neoplastic lesions [42]. Notably, a hypoxia-associated gene program is already active at the metaplastic stage. Hypoxia induces reactive oxygen species (ROS)[43], likely explaining the enrichment of the ROS pathway across all three species. Consistently, we observed a pronounced increase in the expression of NFE2L2, the master regulator of oxidative stress responses, indicating metabolic adaptation to elevated ROS. Despite the hypoxic environment, pancreatic tumour cells still rely on oxidative phosphorylation (OXPHOS) [44], unlike many other tumours that predominantly use glycolysis. Accordingly, the OXPHOS signature is observed in all three species, whereas the glycolysis signature emerges only in zebrafish and human metaplastic cells.

Using additional MSigDB gene signature sets to refine our GSEA analysis, we identified a significant enrichment of the “Gruetzmann pancreatic cancer UP” signature[45] in all three species (Figure 5E). This signature was originally derived from a meta-analysis aimed at identifying consistently differentially expressed genes in pancreatic ductal adenocarcinoma (PDAC) across four independent gene expression datasets. The enrichment of the Gruetzmann signature indicates that a substantial number of markers typically associated with PDAC are already present at metaplastic stages, suggesting that molecular hallmarks of pancreatic cancer emerge at early stages of tumourigenesis, long before the onset of invasive disease.

Overall, these results show that the initiation and progression of pancreatic tumourigenesis involve a set of pathways shared across distantly related species, revealing a conserved mechanism of tumourigenesis.

### Metaplastic cells re-activate a broad spectrum of genes belonging to the pancreatic developmental program

During the transition from acinar to ductal-like cells, studies in mammals have shown that acinar cells acquire certain embryonic progenitor characteristics by expressing several key transcription factors critical for pancreatic development and differentiation (reviewed in[8]). Notably, GATA6, HNF1B, SOX9, SOX4, PDX1 and ONECUT1 have been reported to be upregulated during metaplasia in mammals.

To assess the extent to which metaplastic cells acquire pancreatic progenitor characteristics, we compared their transcriptomic profile with that of multipotent pancreatic progenitor cells, which we transcriptionally profiled by RNA-seq in this study. In zebrafish, these progenitor cells emerge around 32 hours post-fertilization (hpf) and can be readily visualized using the Tg(*ptf1a*:eGFP)^jh1^ transgenic line (Figure 6A). Following microdissection of the pancreatic region, this reporter line enabled us to isolate multipotent pancreatic progenitor cells at 38 hpf by fluorescence-activated cell sorting (FACS). RNA sequencing was subsequently performed on GFP-positive (GFP⁺) and GFP-negative (GFP⁻) populations, revealing 1,411 genes upregulated in the *ptf1a*:GFP⁺ cells (Log2FC > 1, padj < 0.05) (Figure 6B, Additional file 2). Among these upregulated genes, several known pancreatic progenitor markers were identified, including *ptf1a*, *pdx1*, *nkx6.1*, *gata6*, and *sox9b*. Interestingly, within the top five upregulated genes, we identified two novel pancreatic progenitor markers, *olfm4.1* and *olfm4.2* (log2FC = 13 and 11.2, respectively). Visible and fluorescent whole-mount in situ hybridization (WISH) were then performed to confirm the validity of this developmental signature. Visible WISH showed strong and spatially restricted expression of olfm4.1 and apoda.2 within the pancreatic region whereas *cxcl14* and *si:dkey-153k10.9* show additional sites of expression (Figure 6C). Fluorescent WISH confirmed that *olfm4.1*, *elf3,* and *nupr1* are expressed within the *ptf1a^+^* multipotent pancreatic progenitor domain (Figure 6D); only *olfm4.1* is confined to this domain, while *elf3* and *nupr1* show broader expression. Collectively, these data validate the accuracy of our signature for multipotent pancreatic progenitors.

**Figure 6.**
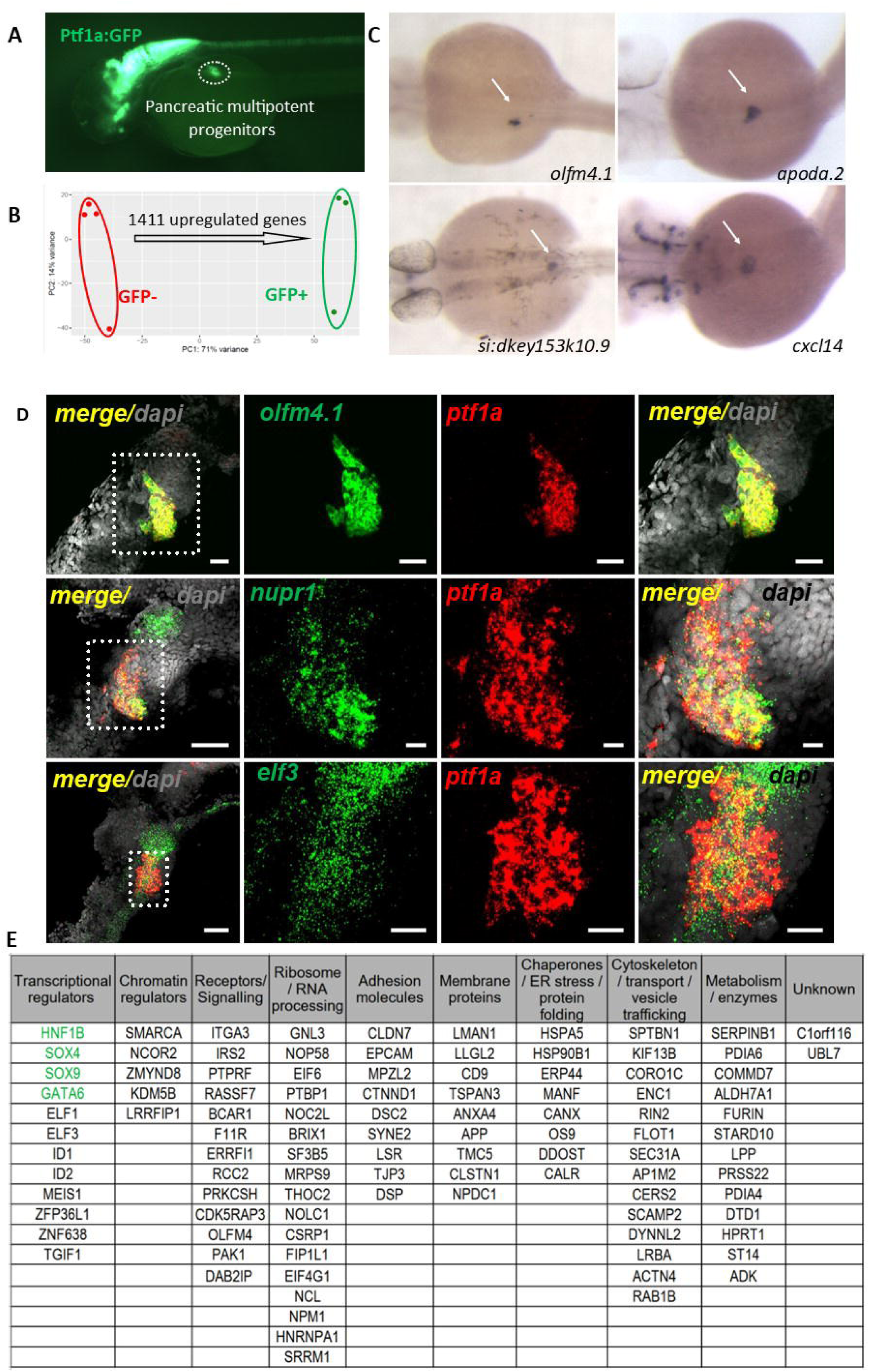
Reactivation of the pancreatic developmental program in metaplastic cells. (A) Visualization of pancreatic multipotent progenitors in a *Ptf1a:GFP* transgenic embryo at 38 hpf. (B) Principal component analysis (PCA) of *rlog*-transformed expression values (DESeq2) for GFP⁺ (green) and GFP⁻ (blue) cells, showing strong correlation among biological replicates. (C) Representative WISH for *olfm4.1*, *apoda.2*, *si:dkey153k10.9*, and *cxcl14* at 38 hpf, showing their expression in multipotent pancreatic progenitors (indicated by white arrows). (D) Double Fluorescent WISH at 35 hpf with olfm4.1, nupr1 and Elf3 revealed in red and ptf1a in green. The region outlined in the left panel is shown in the right panels. Scale bars: 20 µm (E) Table summarizing the main function of the 102 developmental genes expressed in multipotent pancreatic progenitors at 38 hpf and upregulated in metaplastic versus acinar cells across zebrafish, mouse, and human; Genes in green have been previously shown to be involved in metaplasia.

The zebrafish developmental multipotent pancreatic signature was converted into 1,151 human orthologous genes. Comparison with the 1,357 genes upregulated in metaplastic cells across the three species revealed 102 commonly enriched developmental genes. A hypergeometric test confirmed that the presence of this conserved developmental transcriptional program is statistically significant (p = 7.12 × 10⁻¹). Among this set of 102 genes, we found the transcription factors *HNF1B*, *GATA6*, *SOX4*, and *SOX9,* which are already known to be involved in metaplasia [8]. This set also includes several transcriptional regulators that have not yet been implicated in metaplasia (Figure 6E). Notably, ID1 and ID2 (Inhibitor of DNA Binding/Differentiation proteins) are predominantly expressed in progenitor cells during development, where they prevent premature differentiation and promote cell cycle progression [46]. Among these, ID2 has been specifically shown to regulate progenitor expansion during pancreatic organogenesis [47] suggesting that its overexpression in metaplastic cells could similarly promote a progenitor-like state. This list also includes the ETS transcription factors Elf1 and Elf3 whose roles in TGF-β and DCLK1/JAK/STAT signalling pathways, [48,49], suggest potential involvement in metaplasia. Besides transcriptional regulators, this set also contains several epigenetic factors like SMARCA, NCOR2, ZMYND8, KDM5B and LRRFIP1 suggesting that chromatin remodelling could a key component of the metaplastic process.

In conclusion, our analysis reinforces the notion that reactivation of pancreatic developmental program is a central feature of the metaplastic transition.

### Trajectory Analysis Identifies a Cytoskeletal Remodelling Program Associated with Cancer Progression

To capture the dynamics of disease progression, we used the Monocle 3 package that reconstructs cellular trajectories by repositioning cells based on their gene expression profiles, arranging them along a pseudotime continuum. In the human dataset, the trajectory starting at acinar cells (point 1) progresses through early metaplastic stage reaching a bifurcation within the intermediate metaplastic cluster (point 2) (Figure 7A). From this branch point, one path leads toward late metaplastic cells (point 3) and cancer cells, while the other follows an alternative metaplastic route ending in point 4. In the mouse dataset, the trajectory displays a similar overall structure (Figure 7B). After starting from acinar cells, the trajectory progresses through early and proliferative stages, reach a bifurcation within the intermediate metaplastic clusters (point 2). From this bifurcation, one branch advances toward late metaplastic (point 3), to eventually reach cancer cells while the other diverges toward distinct intermediate metaplastic states (point 4).

**Figure 7:**
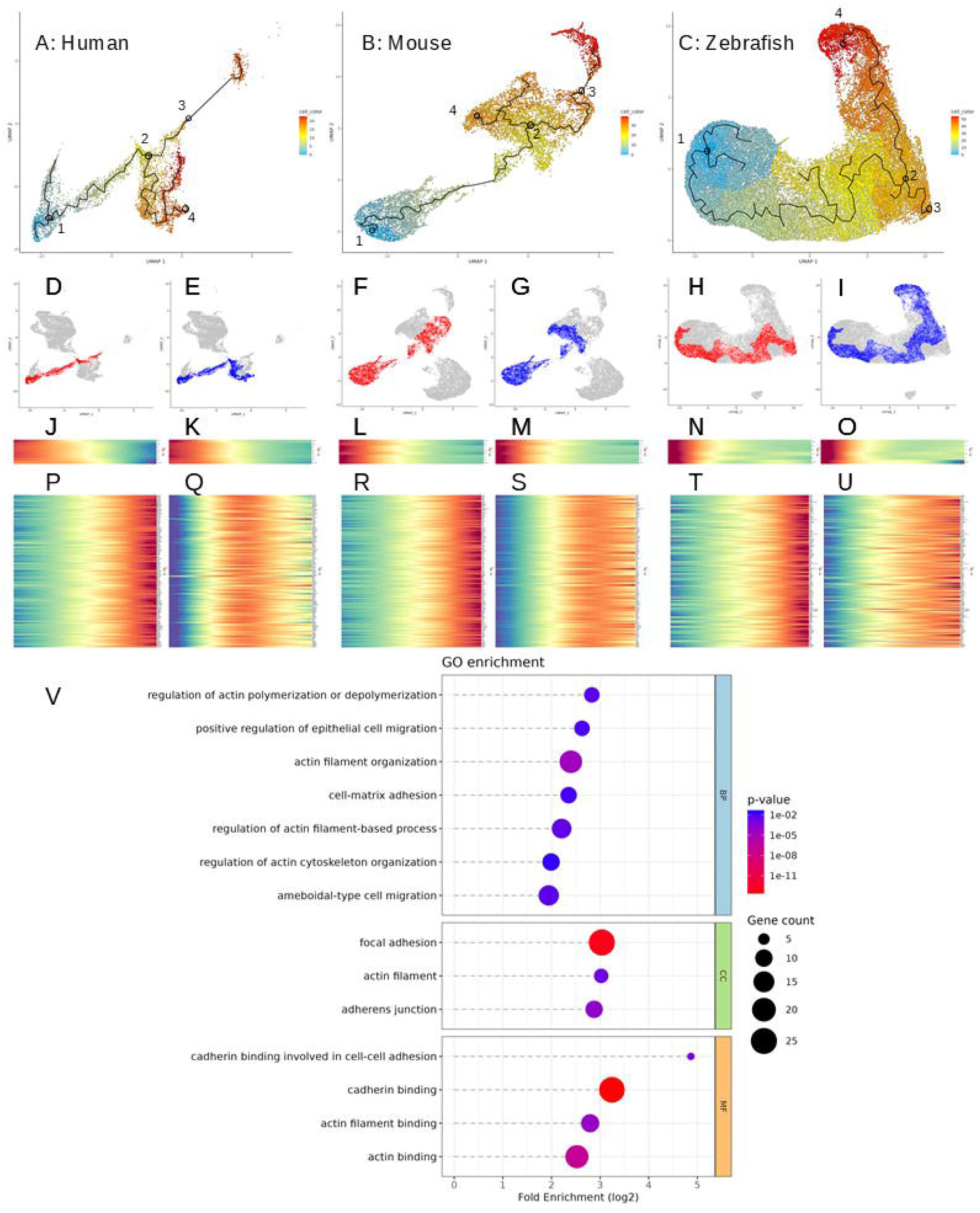
Trajectory Analysis Identifies a Cytoskeletal Remodelling Program in late-metaplastic cells. **A–C**: Representation of the trajectories followed by tumour cells, starting from healthy acinar cells, in human (A), mouse (B), and zebrafish (C). **D–I:** Visualisation of cells included in the late-metaplastic trajectories highlighted in red ( 1->2->3; D,F;H) and the alternative trajectories highlighted in blue (1->2->4; E,G,I). **J-O:** Expression profile of a set of eight acinar markers across cells along both trajectories. **P-U:** Expression profiles of a set of 158 (P-S) or 149 (T-U) genes across cells along both trajectories. Data were z-score normalized independently for each gene and for each plot, such that the displayed values represent relative changes in expression along the trajectory, between the minimum and maximum observed for each gene. **V** : Gene Ontology enrichment analysis of the set of 158 genes. BP, CC and MF represents Biological Pathway, Cellular Component and Molecular Function, respectively.

For both human and mouse dataset, we extracted the cells that form the two main trajectories : cancer trajectory highlighted in red (1->2->3; Figure10 D,F) and the alternative trajectory highlighted in blue (1->2->4; Figure 7 E,G) and analysed their gene expression profiles along these paths. As expected, we observe a significant loss of acinar markers, such as CTRB1, ELA3, and AMY2A along both trajectories (Figure 7, J-M). We then focused on genes selectively upregulated in both species along the branch leading to late metaplastic cells (point 2 → 3), but not along the alternative branch (point 2 → 4). Indeed, these genes may represent key regulators of the metaplasia-to-cancer transition. This analysis revealed 158 genes whose expression increases specifically along the cancer-directed branch in both species, peaking in late metaplastic cells (Figure 7 P,R) while declining along the alternative trajectory (Figure 7 Q,S). Notably, this set includes 13 transcription factors (KLF5, KLF6, FOXQ1, RUNX1, FOSL2, BHLHE40, ETS2, TGIF1, TSHZ2, ZBTB7A, SREBF2, HIF1A and CUX1), suggesting that coordinated activation of specific regulatory modules accompanies the transition from metaplasia to cancer. Gene Ontology enrichment analysis of these 158 genes revealed a strong enrichment for biological processes related to actin dynamics, including “regulation of actin polymerization or depolymerization,” “regulation of actin filament–based processes,” and “regulation of actin cytoskeleton organization,” highlighting extensive remodelling of the cytoskeletal network (Figure 7V). These changes are consistent with dynamic regulation of cell shape and polarity, reflecting increased structural plasticity. The enrichment in *“cell migration”* and *“ameboidal-type cell migration”* suggests the acquisition of a more motile phenotype characterized by actin-driven contractility and transient cell–matrix interactions. The presence of *“focal adhesion”* further supports this interpretation, suggesting activation of integrin-mediated signalling and mechanical coupling between the actin cytoskeleton and the extracellular matrix—processes essential for traction and directional movement. Altogether, these findings suggest that late metaplastic cells have undergone extensive cytoskeletal and adhesive remodelling, acquiring the structural and functional traits associated with increased motility and the potential to become invasive. Intriguingly, we found no molecular evidence supporting a full epithelial-to-mesenchymal transition (EMT), such the induction of the canonical EMT factors (SNAI1/2, TWIST1/2 and Zeb1/2) and the upregulation of vimentin and downregulation of CDH1 expression. These typical EMT markers were also not induced in any of the other metaplastic cells. Instead, the upregulation of VCL (vinculin), which is preferentially associated with remodelling rather than stable adherens junctions, points to a shift from rigid epithelial junctions toward more dynamic and flexible adhesion complexes that may facilitate cell migration while preserving partial intercellular connection [50]. These observations support the new concept that PDAC invasion may occur through alternative migratory behaviors, such as collective or clustered migration, rather than through a fully mesenchymal EMT-driven, single-cell mode of invasion [51,52].

In zebrafish, the main trajectory also progresses from healthy acinar cells through early and intermediate metaplastic states before bifurcating: one branch enters a proliferative program (S phase followed by G2/M), while the other advances toward late metaplastic stages. However, because the six sequenced tumours did not progress to the PDAC stage, no cancer cells were detected in the zebrafish dataset, preventing us from extending the trajectory beyond the late stage and thus from reconstructing a “cancer trajectory.” Nevertheless, most of the 158 genes significantly upregulated in the late metaplastic cluster in mouse and human also exhibit a gradual increase in expression along the late-metaplastic branch in zebrafish (149 out of 158), with the majority reaching peak expression at the terminal end of the trajectory (Figure 7T). However, this upregulation is generally weaker than in mouse and human and is often confined to a small subset of late metaplastic cells at the very end of the trajectory.

These observations indicate that the transcriptional program activated at the terminal stages of metaplasia in zebrafish closely mirrors that observed in mammals.

### Dynamic Transcription Factor Activity Along the Metaplasia–Cancer Trajectory

To further explore which transcriptional programs are active during acinar-to-cancer progression, we analysed regulon activity using the SCENIC pipeline. SCENIC evaluates whether a transcription factor and its predicted target genes are co-expressed in single cells [53]. When such co-expression is observed, the corresponding regulon is defined as active in that cell. This prediction relies on the enrichment of the transcription factor binding motifs in the regulatory regions of the candidate target genes. For mouse and human, two databases are available that report DNA motifs significantly overrepresented in the vicinity of transcription start sites (TSS), either within a 10 kb window around the TSS or within 500 bp upstream of the TSS. Unfortunately, equivalent databases are not yet available for zebrafish.

Using both the 10 kb and 500 bp motif databases (see Methods), we searched for regulons active in acinar and metaplastic cells, with a particular focus on those that become progressively activated along the trajectory leading toward cancer. To that end, human metaplastic cells were further subdivided to better resolve their heterogeneity (Figure 8A, upper panel). Figure 8 A,B presents a binarized heatmap of the regulons identified by SCENIC as active in human and mouse samples.

**Figure 8:**
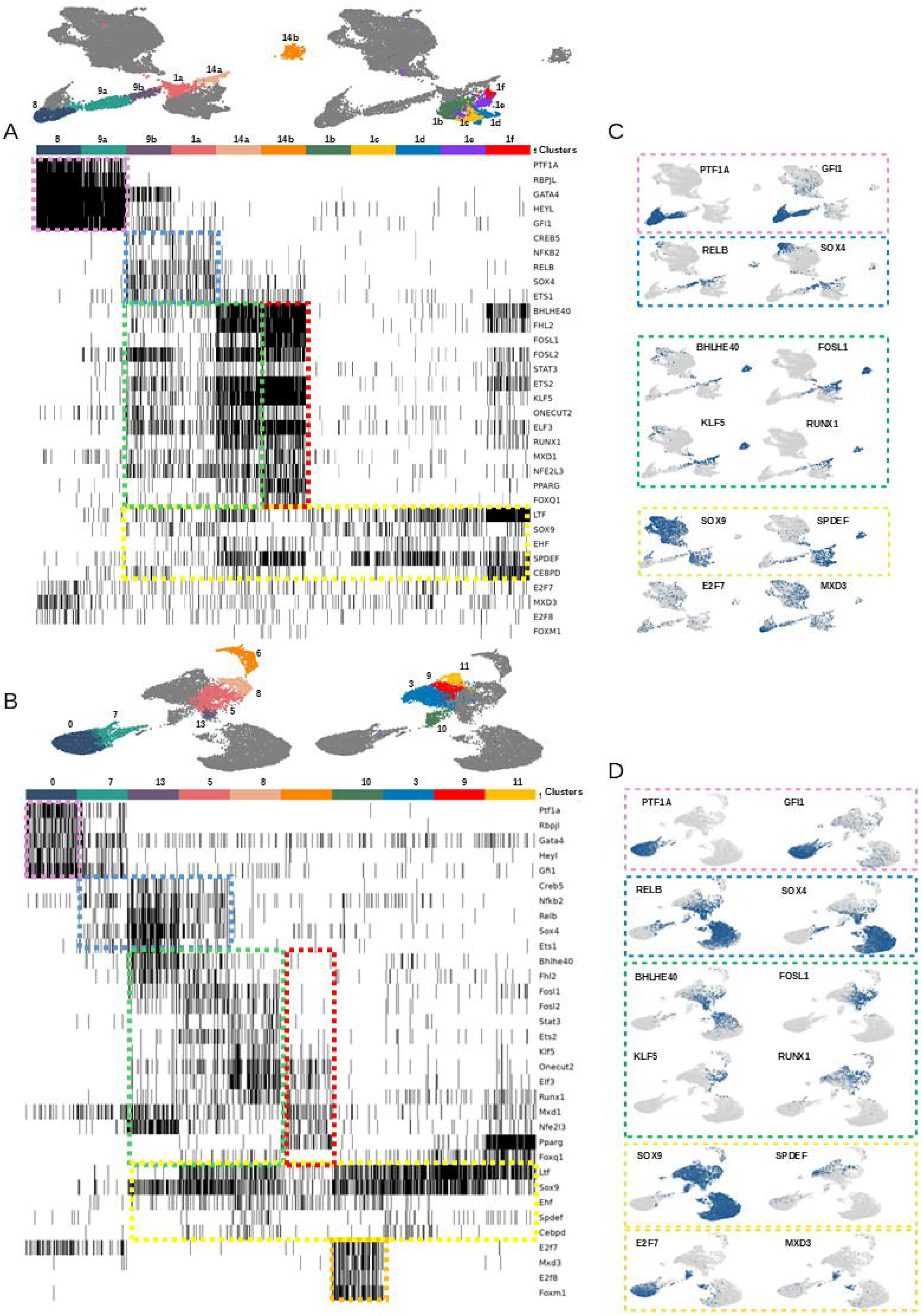
SCENIC-based Identification of Active Transcriptional Regulons. **A–B:** Binarized heatmaps of regulons identified by SCENIC as active in human (A) and mouse (B) samples. **C–D**: Feature plots showing the activity of selected regulons in human (C) and mouse (D) samples.

Human acinar (cluster 8) and acinar-like cells (cluster 9a) displayed strong and specific activity of regulons driven by known acinar regulators (RBPJL, PTF1A, GATA4) and by the Notch effector HeyL [54], thereby validating the SCENIC approach (Figure 8A). A regulon for Gfi1, a factor linked to exocrine cell differentiation, was also detected [55]. The heatmap also revealed regulons that are activated transiently at early stages of metaplasia (highlighted in blue) such as SOX4 and RELB, whose regulon activity feature plots clearly confirm this highly restricted activation (Figure 8C). Interestingly, we identified a set of 14 regulons (in green) that display high—and generally increasing—activity along the trajectory leading to cancer (clusters 9b →1a→14a). Most of these regulons reach maximal activity in the late metaplastic cluster 14a and remain strongly active in the cancer cell cluster 14b (in red). In contrast, their activity is much lower in the other intermediate metaplastic clusters (clusters 1b-1f), with the exception of cluster 1f, (adjacent to cluster 14a), which displays partial regulon activities. Among these 15 regulons are those driven by the transcription factors KLF5, FOXQ1, RUNX1, FOSL2, BHLHE40, and ETS2, which we previously identified in our trajectory analysis as being upregulated in late metaplastic cells. Visualization of their regulon activity using feature plot illustrates this restricted activation along the trajectory (Figure 8C). Finally, we also identified regulons that remain active across all metaplastic cells (shown in yellow), such as the Sox9, LTF and EHF regulons (Figure 8A,C).

SCENIC analysis of the mouse dataset (Figure 8B) also revealed strong acinar regulon activity for PTF1A, RBPJL, HEYL, and GATA4, although GATA4 is less acinar-specific than in human. Early metaplasia regulons, including SOX4, RELB, and CREB5, were similarly activated (shown in blue). Nine of the 14 regulons enriched along the human acinar-to-cancer trajectory were also selectively activated along the mouse trajectory, including BHLHE40 and KLF5. The remaining five, such as RUNX1, FOXQ1, and PPARG, showed broader activity across all metaplastic cells. Only six of the 14 regulons remained active in mouse cancer cells, compared with humans. As in human, SOX9, LTF and EHF were active across all metaplastic cells (in yellow). As for the proliferative cluster, unique to mouse and located along the cancer trajectory, it exhibits high activity of cell cycle–associated regulons (E2F7, E2F8, MXD3, FOXM1, highlighted in orange) but minimal or no activity of other metaplastic regulons, except for SOX9, EHF and LTF. Visualisation of selected regulons for each category is shown in Figure11C-D.

Together, these analyses reveal a conserved set of regulons, highlighted in green that is shared across the mouse and human cancer trajectories and becomes progressively activated as cells advance toward malignancy. In contrast, other regulons—such as the SOX9 regulon highlighted in yellow — display widespread activity across all metaplastic cell states, acting as key regulators of metaplasia rather than indicators of progression toward cancer. This distinction underscores the existence of both progression-associated and state-defining regulatory programs within the metaplasia-to-cancer continuum.

## Discussion

In this study, we generated a robust and highly efficient zebrafish model of pancreatic tumourigenesis by expressing Kras^G12D^ **i**n mature acinar cells using the GAL4/UAS system. Oncogenic Kras activation leads to the development of pancreatic tumours, a process that is markedly accelerated by loss of p53 function. Histological and molecular analyses of the p53^m/m^ tumours show that all tumours undergo a progressive acinar-to-ductal metaplastic transition, characterized by loss of acinar identity and induction of ductal markers such as *sox9b*. While most tumours remain at a metaplastic stage, a subset acquires invasive properties and displays key hallmarks of human PDAC, including glandular differentiation and desmoplasia. Single-cell transcriptomic analyses reveal that metaplasia encompasses a continuum of cellular states, with cells gradually evolving through a range of intermediate states, rather than undergoing an abrupt state transition. Cross-species comparisons with mouse and human scRNAseq datasets uncover a highly conserved transcriptional landscape underlying metaplasia, indicating that early molecular features of pancreatic cancer are shared across vertebrates. In all three species, this metaplasia process reactivates a whole series of factors normally expressed in multipotent pancreatic progenitors, reinforcing the notion that the reactivation of developmental processes is an important aspect of metaplasia. High concordance across species is also observed when reconstructing tumour cell trajectories from acinar to cancerous states, revealing shared gene expression changes along this progression. Notably, these analyses identified a set of genes involved in cytoskeletal remodelling and cell migration, which are specifically upregulated in the late metaplastic stage, just prior to the transition to the cancerous state. Among these genes are several transcription factors, such as KLF5, FOXQ1, RUNX1, FOSL2, BHLHE40, and ETS2. Importantly, SCENIC analyses revealed that these factors display high activity in this transition zone, suggestive of a key role in the progression from metaplastic to cancerous cells. Altogether, these findings position acinar-to-ductal metaplasia as an important and conserved step in pancreatic cancer initiation and progression and establish zebrafish as a powerful model to dissect early events driving PDAC.

### A robust zebrafish model with transient Kras expression recapitulates early pancreatic tumourigenesis

Compared to the two previously described zebrafish acinar-derived pancreatic tumour models [24,25], our system is considerably more efficient and robust. All p53m/m fish develop tumours by one year of age while tumour incidence is only 11% in the model (ptf1a-Cre^ERT2^; ubb:LSL:GAL4-VP16; UAS:eGFP-KRAS^G12V^) described by Park and Leach [24] and 40% in the model (Ela3l:CRE; ubb:Lox-Cherry-Lox-GFP:KRAS^G12D^) developed by Oh and Park [25]. This lower incidence may be explained, at least in part, by the use of a wild-type genetic background, whereas our data demonstrate that p53 mutation dramatically accelerates tumour onset and progression. Intriguingly, the tumours characterized by Oh and Park closely resembled pancreatic endocrine tumours. In contrast, we never detected expression of endocrine hormones in either our p53^m/m^ or p53^+/+^ tumours (data not shown). Like the p53^m/m^, the p53^+/+^ tumours display metaplastic features without any sign of endocrine differentiation. The reason for this difference remains unknown. It is also important to note that tumour onset occurs much earlier than suggested by the tumour incidence curve (Figure 1H). This curve was generated based on visual detection of tumours in living fish, which only allows identification once tumours are large enough to cause a noticeable protrusion. Histological analysis of a random subset of fish dissected at 5 months revealed that all of them already harboured tumours (n=7), not yet visible to the naked eye, demonstrating that tumours arise substantially earlier than anticipated (data not shown). The model used here, in which KalTA4 expression is driven by the *ela3l* promoter, is characterized by transient Kras^G12D^ expression. Indeed, *ela3l* is the most strongly downregulated acinar transcripts during acinar-to-ductal metaplasia, likely reflecting loss of *ela3l* promoter activity during this process. This in turn silences KalTA4, thereby extinguishing GFP–Kras^G12D^ expression. Consistent with this, 36% of tumours display a complete or near-complete loss of GFP signal (Supplementary Figure 1). The silencing of Kras^G12D^ is likely the reason why a relatively limited number of tumours progress beyond the metaplasia stage. Nevertheless, a small subset does reach the PDAC stage (approximately 7%, Table 1A), and this percentage increase drastically when considering only tumours with a complete or near-complete loss of GFP fluorescence (57%, 8 out of 14, Table 1B). This suggests that the PDAC stage can be reached even when Kras is no longer active, consistent with prior studies in mice [56]. Indeed, using a doxycycline-inducible Kras^G12D^-driven PDAC model, Kapoor et al. show that doxycycline withdrawal initially leads to tumour regression. Relapse tumours subsequently emerge, with approximately half harbouring an activated Yap1/Tead2 transcriptional program enabling Kras-independent proliferation. In this context, **s**ingle-cell RNA sequencing of the zebrafish PDACs would be particularly informative for identifying factors or pathways that compensate for the loss of Kras activity. Such factors are likely to represent key targets in the context of anti-Kras therapies currently under development [57]. Indeed, following an initial period of remission, tumours are likely to acquire resistance to Kras inhibition by engaging Kras-independent pathways.

### Conserved transcriptional programs and signalling pathways in pancreatic metaplasia

Cross-species comparison of the metaplastic process in zebrafish, mouse, and human revealed that this process is highly conserved and allows the identification of transcriptomic changes that are likely essential for disease progression. Among these conserved factors are a series of transcription factors expressed in multipotent pancreatic progenitors, including HNF1B, SOX9, SOX4, and GATA6, which are already known to play key roles in metaplasia[8]. We also identified novel developmental regulators such as *olfm4* that displays an early, restricted and strong expression in zebrafish multipotent progenitor cells as demonstrated by *in situ* hybridization (Figure 6C-D) suggesting a developmental role in this organ. At later stages, *olfm4* is also expressed in the mature intestinal crypts, where it is a well-established marker of LGR5⁺ intestinal stem cells [58]. Interestingly, OLFM4 has also been identified as a biomarker of gastric intestinal metaplasia (IM) [59], a condition characterized by the replacement of normal gastric mucosa with intestinal-type epithelium in response to chronic gastric inflammation. Similar to pancreatic metaplasia, IM represents an intermediate stage in the progression from premalignant lesions to malignant transformation in gastric cancer. Notably, OLFM4 was shown to promote the progression of intestinal metaplasia through activation of the MYH9/GSK3β/β-catenin signalling pathway [60]. Taken together, these findings strongly suggest that OLFM4 may also play an important role in pancreatic metaplasia, a possibility that warrants further investigation. The induction of OLFM4 could be due to the TNF-α/NF-κB signalling pathway which**_**is activated in all three analysed**_**species (Figure 5E). Indeed, it has been shown that the regulation of OLFM4 expression in myeloid precursor cells relies on NF-κB transcription factor [61]. In addition, in intestinal epithelial cells, TNF-α, in synergy with Notch signalling, can strongly induce OLFM4 expression [62]. Although Notch pathway enrichment was not identified as significant in the GSEA analysis, upregulation of its key downstream effector, HES1, is observed across all three species, suggesting a Notch activity in metaplastic cells.

The developmental transcription factor ELF3 may also represent an important regulator of metaplasia, as it has been identified as a potent activator of TGF-β signalling in the intestine, primarily through binding to two adjacent ELF3 sites in the promoter region of the type II TGF-β receptor gene (*T*β*R-II*) [48,63] The upregulation of TGF-β signalling, observed across the three species (Figure 5E) has been shown to drive metaplasia progression [42]. However, this activation is likely transient, as a frequent loss of the tumour suppressor *SMAD4*, a key downstream effector of TGF-β signalling is often observed at advanced PDAC stages.

### Cross-species trajectory analysis identifies conserved regulators of metaplasia-to-cancer progression

High concordance across species is also observed in reconstructed tumour cell trajectories. In both human and mouse datasets, trajectories originating from acinar cells progress through an early metaplastic stage and reach a bifurcation within the intermediate metaplastic cluster. From this branch point, one trajectory leads toward late metaplastic and cancer cells, whereas the other diverges toward distinct intermediate metaplastic states (Figure 7A–B). Interestingly, we identified a set of 158 genes that are selectively upregulated along the branch leading to cancer cells. These genes are primarily involved in actin cytoskeleton dynamics and focal adhesion remodelling, suggesting increased structural plasticity and motility. However, we did not observe along this branch induction of the canonical EMT program, including the EMT-associated transcription factors SNAI1, SNAI2, TWIST1/2, ZEB1, and ZEB2, nor the typical EMT marker switch characterized by vimentin upregulation and CDH1 downregulation. This suggests that late metaplastic cells do not undergo a classical EMT program. This is consistent with previous reports showing that combined genetic deletion of the key EMT transcription factors *Snai1* and *Twist1* in mouse KPC models does not impair the emergence of invasive PDAC, nor prevent systemic dissemination or metastasis [64]. Notably, these knockout cells retain an epithelial phenotype and display enhanced collective migration *in vitro*. In agreement with this, three-dimensional histological analyses of PDAC have shown that collective migration represents the predominant mode of invasion, whereas single-cell invasion is exceedingly rare [65]. Nevertheless, as single-cell RNA-sequencing analyses have revealed that pancreatic cancer cells are distributed along an epithelial–mesenchymal continuum in both human and murine tumours [64], multiple invasion strategies likely coexist. In this context, mesenchymal-like cells may be more prone to disseminate as single cancer cells, whereas epithelial-like cells that retain cell–cell adhesions may preferentially disseminate as multicellular clusters. Altogether, these observations support the emerging concept that tumour cells exhibit remarkable plasticity in their invasive programs and can dynamically switch between distinct modes of invasion to adapt to different microenvironments [52].

Beyond cytoskeletal regulators, this 158-gene signature includes thirteen transcription factors, among which KLF5, FOXQ1, RUNX1, FOSL2, BHLHE40, and ETS2 display high and specific regulon activity within the late-metaplastic zone (Figure 8), suggesting a potential role in the progression from metaplastic to cancerous states. Among these factors, KLF5 appears particularly relevant, as it has been reported in nasopharyngeal carcinoma to control a transcriptional program that includes α-actinin-4 (ACTN4). This program influences actin cytoskeleton remodelling, enhances cell motility, and promotes metastatic dissemination [66]. The roles of FOSL2, BHLHE40, and ETS2 in the metaplasia–cancer transition appear particularly worthy of investigation, as to date very limited data are available regarding these factors in pancreatic cancer.

Importantly, this cross-species conservation of the tumour cell trajectory also extends to zebrafish. The branch leading to the late metaplastic stage exhibited upregulation of the majority of the 158-gene signature, despite the fact that a bona fide cancer trajectory could not be defined because no overt cancer cells were sequenced. These observations indicate that the transcriptional program activated at the terminal stages of metaplasia in zebrafish closely mirrors that observed in mammals. However, this upregulation was generally weaker that what was observed in mice and human and was often detected in a small subset of late metaplastic cells at the very end of the trajectory. This is notably the case for *klf5l, anxa1a* and *onecut2.* Therefore, it would be interesting to determine whether overexpression of these factors, either individually or in combination, is sufficient to drive progression toward more advanced PDAC-like stages.

The identification of at least two distinct trajectories—one leading to cancer and the other to different metaplastic subsets—raises the question of whether the alternative path represents a biological dead end. A similar issue was previously raised by Schlesinger et al. [11]who showed that rather than a continuous progression from acinar cells to metaplastic cells and then to cancer cells, Monocle analyses indicated that acinar cells could follow two divergent trajectories, one leading to stomach-like metaplastic cells and the other to cancerous states. Determining whether these alternative trajectories truly represent “dead ends” remains challenging. It is likely that multiple trajectories coexist and that, depending on various parameters—such as the tumour microenvironment, hypoxia, KRAS dependency, epigenetic state, and metabolic constraints- the routes leading to PDAC may differ. In this context, a path that appears to be a dead end in one model may not be so in another. Elucidating the full spectrum of possible trajectories is therefore an important step from a therapeutic perspective, as blocking one path may push tumour cells to progress through alternative routes.

### Dynamic transcription factor regulon activity during metaplasia-to-cancer transition

To identify the regulatory networks important for pancreatic tumourigenesis, we measured regulon activity using SCENIC, which infers whether a transcription factor is functionally active based on the co-expression of its target genes. This often provides a more accurate measure of functional activity of a transcriptional factor than its mRNA level. This analysis allowed us to identify a set of 14 regulons whose activity progressively increases along the trajectory toward cancer, reaching maximal levels in late metaplastic and cancer cells. Visualization of their activity clearly shows that, for example, FOSL2, KLF5, and RUNX1 are specifically active along this trajectory (Figure 8C), highlighting a regulatory program in these cells that is clearly distinct from the alternative trajectory. At the same time, we also observed that some regulons are active across all metaplastic cells. This is notably the case for SOX9, a well-established marker of metaplastic cells, whose role is to repress acinar genes and activate ductal gene programs [29]. Other regulons appear to be activated at very early stages of the process, such as SOX4, which is among the most highly up-regulated genes during ADM and has been implicated in the regulation of cellular dedifferentiation [67]. The role of additional early-active regulons, such as CREB5, RELB, and NFKB2, would also merit further investigation.

### Identification of early biomarkers of PDAC

In this study, we characterized in detail the early stages of pancreatic tumourigenesis, which will help guide the identification of early biomarkers. Nevertheless, their discovery remains challenging as in addition to oncogene-driven ADM, acinar cells can also undergo spontaneous ADM in the context of pancreatic inflammation [8,68]. This inflammatory-associated metaplasia represents a physiological and reversible process that contributes to acinar cell regeneration following tissue injury. This overlap between physiological and oncogenic metaplasia complicates the identification of *bona fide* early PDAC markers, as many features of ADM are shared between these two contexts. Therefore, the discovery of early diagnostic markers will require the identification of genes that are selectively associated with oncogenic metaplasia, but absent from inflammatory, regenerative ADM. In this regard, OLFM4 emerges as a particularly promising candidate. Our data indicate that within human datasets, OLFM4 is expressed in oncogenic metaplastic cells but not in the inflammatory metaplastic cells. Moreover, OLFM4 is a secreted protein that remains expressed in established PDAC, making it attractive from a translational perspective. Nevertheless, OLFM4 is not pancreas-specific and has previously been reported as a biomarker in other gastrointestinal malignancies [69]. Future work will therefore be required to perform a more comprehensive analysis aimed at identifying additional markers that, alone or in combination, could improve the specificity and robustness of early PDAC detection.

## Conclusions

This study reveals that the molecular programs driving the earliest steps of pancreatic ductal adenocarcinoma development are highly conserved across species. Using a zebrafish model of oncogenic KRAS–induced acinar-to-ductal metaplasia combined with single-cell transcriptomic and trajectory analyses, we identify conserved developmental and regulatory networks that are reactivated during preinvasive disease. Importantly, we uncover a conserved transcriptional transition characterized by the induction of cytoskeletal and migratory gene programs, marking the shift from metaplasia to malignant progression. These findings provide a framework for identifying early biomarkers of PDAC and define evolutionarily conserved pathways that represent promising targets for preventive and early-intervention strategies.

## Supporting information

additionnal file 1

additionnal file 2

## List of abbreviations

PDAC: Pancreatic ductal adenocarcinoma
Hpf: hours post fertilization
Dpf: days post fertilization
ADM: acinar to ductal metaplasia
PanIN: Pancreatic Intraepithelial Neoplasia
IPMN: Intraductal Papillary Mucinous Neoplasm
scRNA-seq: single-cell RNA sequencing
HE: haematoxylin and eosin
AB: Alcian Blue
WISH: whole-mount *in situ* hybridization
DEG: differentially expressed genes
S.E: standard errors
F.C: Fold change
FDR: false discovery rate
N.D.: not done
Wt: wild-type

## Declarations

### Ethics approval

The fish were maintained according to national guidelines and all experiments described were approved by the Ethics Committee of the University of Liège (protocol numbers 16-1851 and 21-2355).

As tumour onset in our transgenic line is unpredictable as tumours can arise at any time between 3 and 12 months of age, animals were euthanized as soon as tumours became visible to the naked eye, in accordance with ethical committee approval. Animal welfare was assessed at regular intervals, and no signs of distress, impaired mobility, or compromised health were observed.

### Consent for publication

Not applicable

### Availability of data and materials

The zebrafish sequencing data (FASTQ files) have been deposited in the European Nucleotide Archive (ENA) at EMBL-EBI under accession number PRJEB107715 and will be made publicly available upon publication (https://www.ebi.ac.uk/ena/browser/view/PRJEB107715). In addition, to maximize data accessibility and impact, a publicly available web-based platform is currently under development, which will enable researchers to easily visualize gene expression profiles across the three species, thereby enhancing the visibility and usability of this resource.

### Authors’ contributions

M.K generated the acinar-derived model. C.G. performed the experiments for Figures 1–6 and Supplementary Figures 1–4, with assistance from E.D. and prepared the corresponding figures. C.L. performed the bioinformatic analyses for Figures 7–11 and Supplementary Figures 5–10 and prepared the figures. A.R.L.P. performed the RNA-seq analyses of *ptf1a:GFP* and conducted the zebrafish visible and fluorescent WISH experiments, and prepared Figure 6. N.B. analysed the histological H&E staining and classified the PDAC samples. C.G., L.F., B.P., and M.V. wrote the main manuscript text. All authors reviewed and approved the manuscript

## Acknowledgements

We thank L. Zon for the tp53^zdf^ line [27] and M. Parsons for the Tg(ptf1a:eGFP)^jh1^ line [71]. We also acknowledge F. Argenton for providing the tg(UAS-e1b:GFP-Kras^G12D^) transgene [21] and M. Distel for the kalTA4 fusion cDNA [26]. We thank the following technical platforms: GIGA-Zebrafish, GIGA-Cell Imaging and Flow Cytometry, the GIGA-Genotranscriptomic, the GIGA-bioinformatics and the GIGA-Immunohistology platforms. We also thank the Service d’Anatomie et Cytologie Pathologiques of Professor Philippe Delvenne, and more particularly the anatopathologists Noëlla Blétard and Claudia Pop, for their careful examination of the zebrafish tumours. We thank also Yves Herremans and the VSTA core facility at VUB (https://vsta.research.vub.be) for the RNAscope. We are indebted to Virginie Vonberg for technical help.

## Supplementary Figures

**Supplementary Figure 1:**
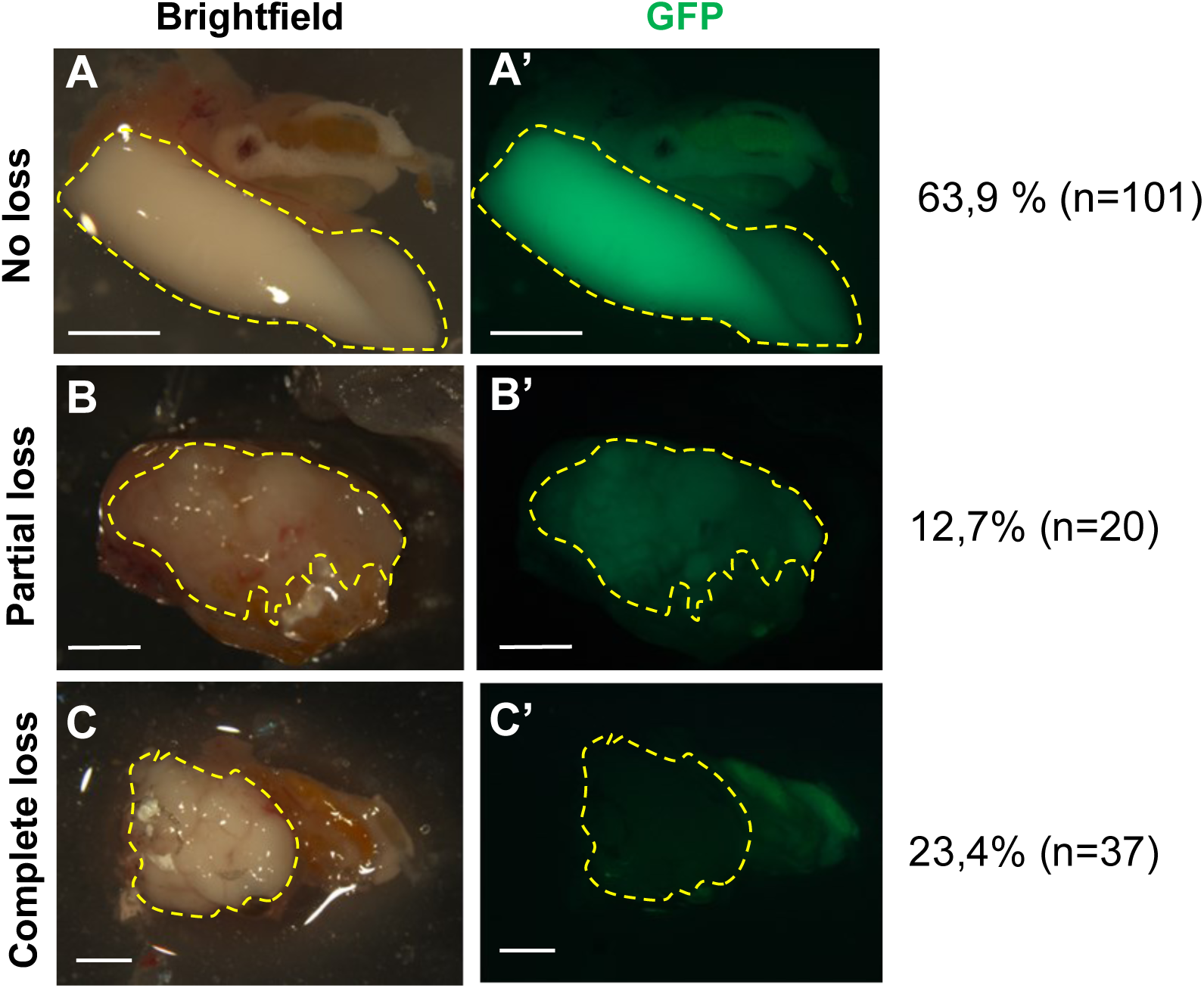
Heterogeneous reduction of GFP-KRAS^G12D^ expression in Ac-K p53m/m tumours. Representative examples of tumours showing no apparent loss (A), partial loss (B) or complete loss (C) of GFP-Kras^G12D^ expression. The percentage and number of fish in each category are indicated on the right. Tumours are outlined with yellow dashed lines. Scale bars=2mm.

**Supplementary Figure 2:**
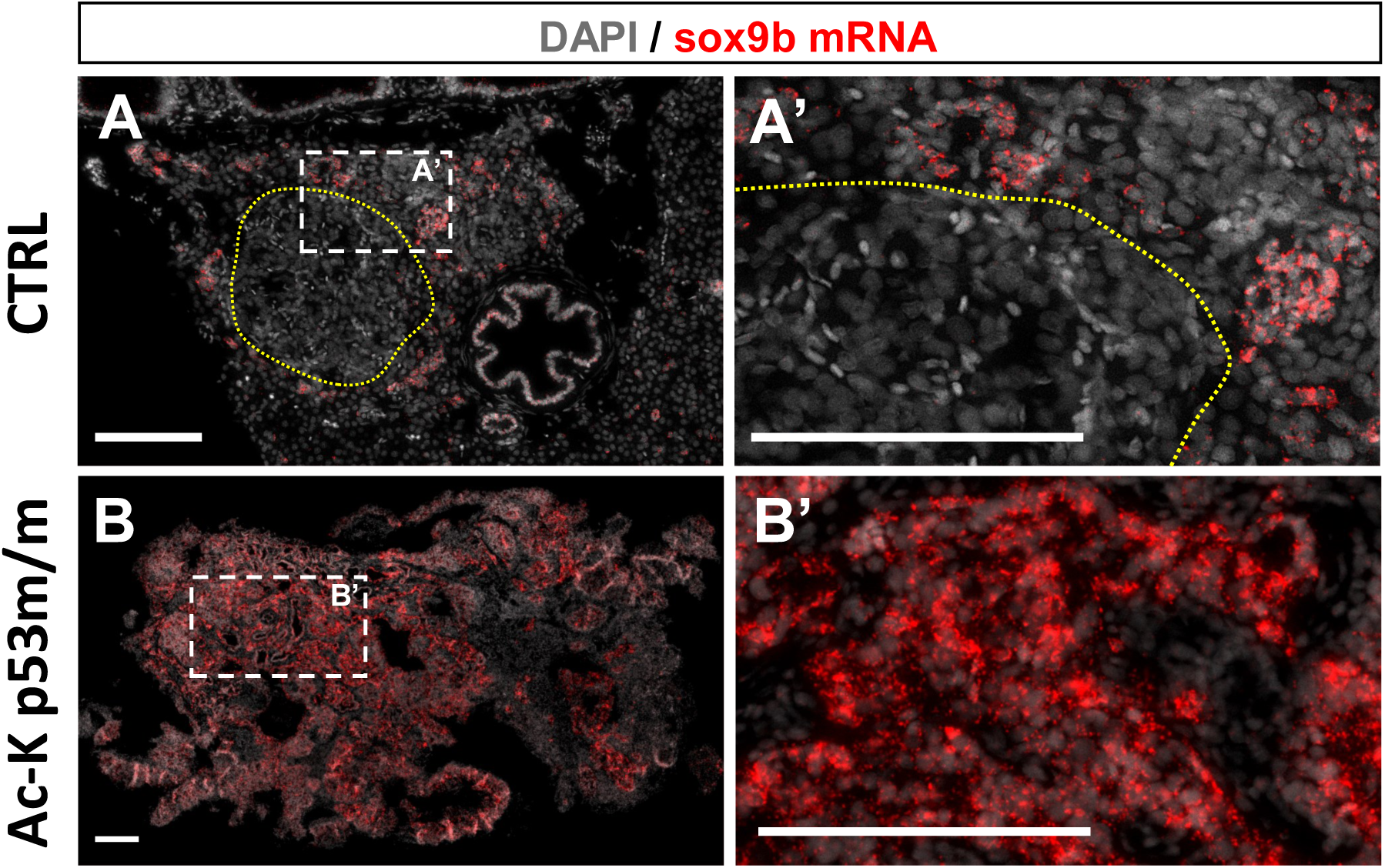
*sox9b* metaplastic marker expression is increased in Ac-K p53^m/m^ tumours. **A-B:** RNAscope detection of *sox9b* mRNA (red) and DAPI nuclear staining (grey) in Ac-K p53^m/m^ tumours and control pancreas. *sox9b* mRNA is ectopically expressed in the tumoral pancreatic tissue (B, B’), whereas in controls *sox9b* expression is restricted to pancreatic ducts (A, A’; arrowheads). The principal islet (i) is outlined with yellow dashed lines. Data are representative of two metaplastic p53^m/m^ tumours and three control pancreata (1 p53^+/+^ and 2 p53^m/m^).

**Supplementary Figure 3 :**
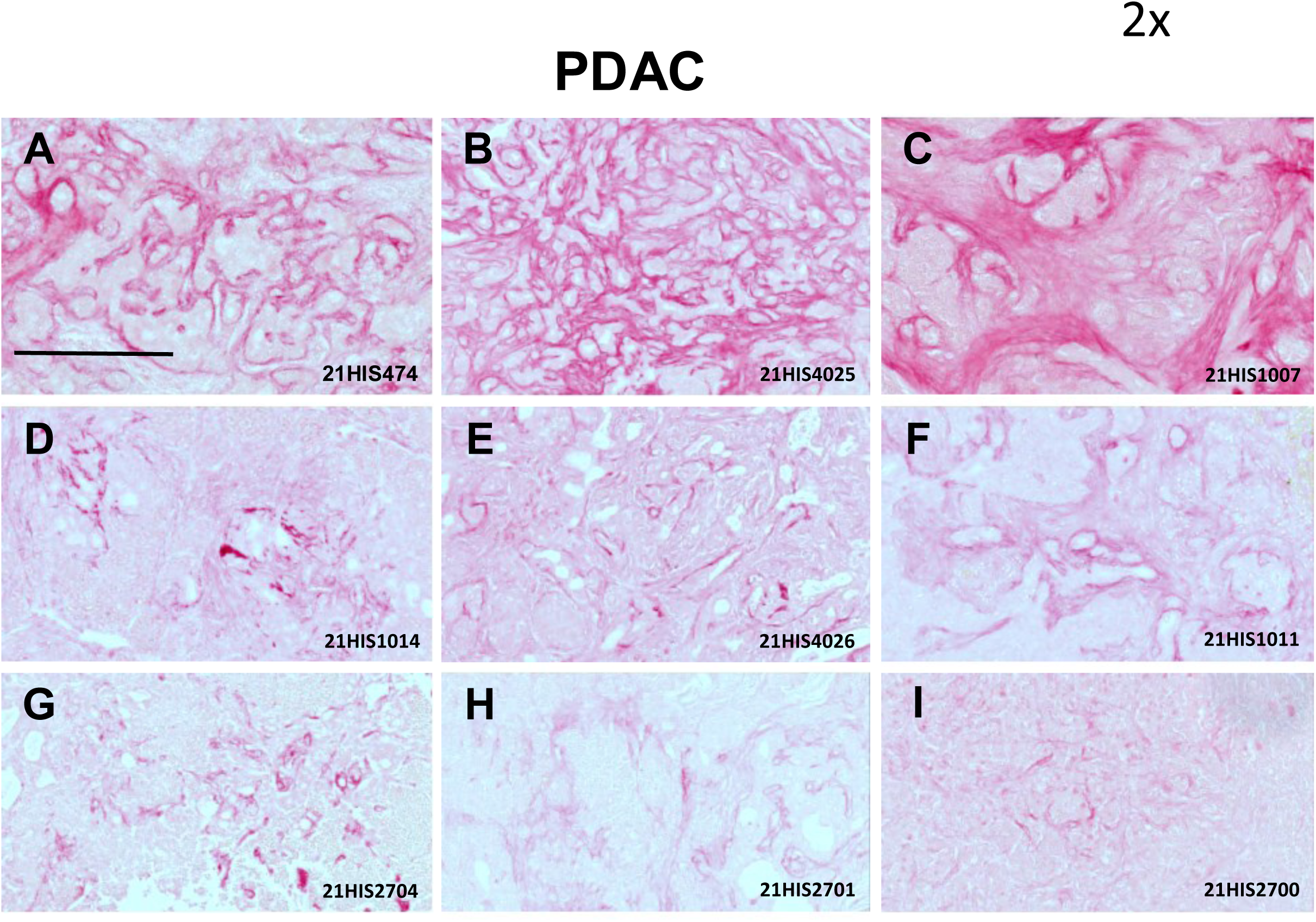
Sirius Red stainings of PDAC. All PDAC samples displayed desmoplasia, with either high (**A-C**) or intermediate (**D-I**) staining levels. Scale bar: 200uµm

**Supplementary Figure 4 :**
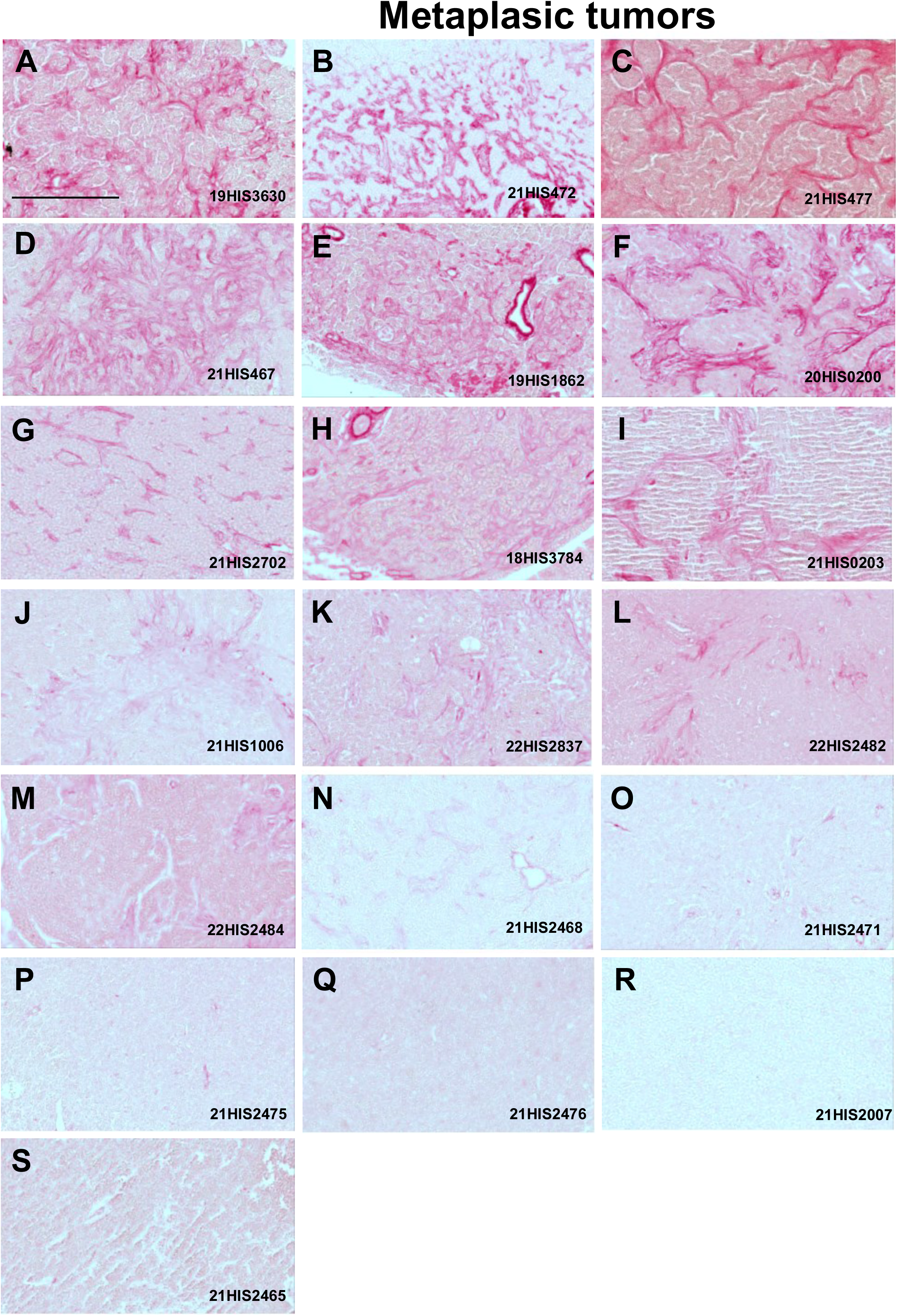
Desmoplasia is detected in several metaplastic tumours. Six samples show high levels of Sirius red staining (A-F), seven tumours exhibited intermediate levels (G–M), whereas low or no detectable staining was observed in six other samples (N–S). Scale bar: 200uµm

**Supplementary Figure 5 :**
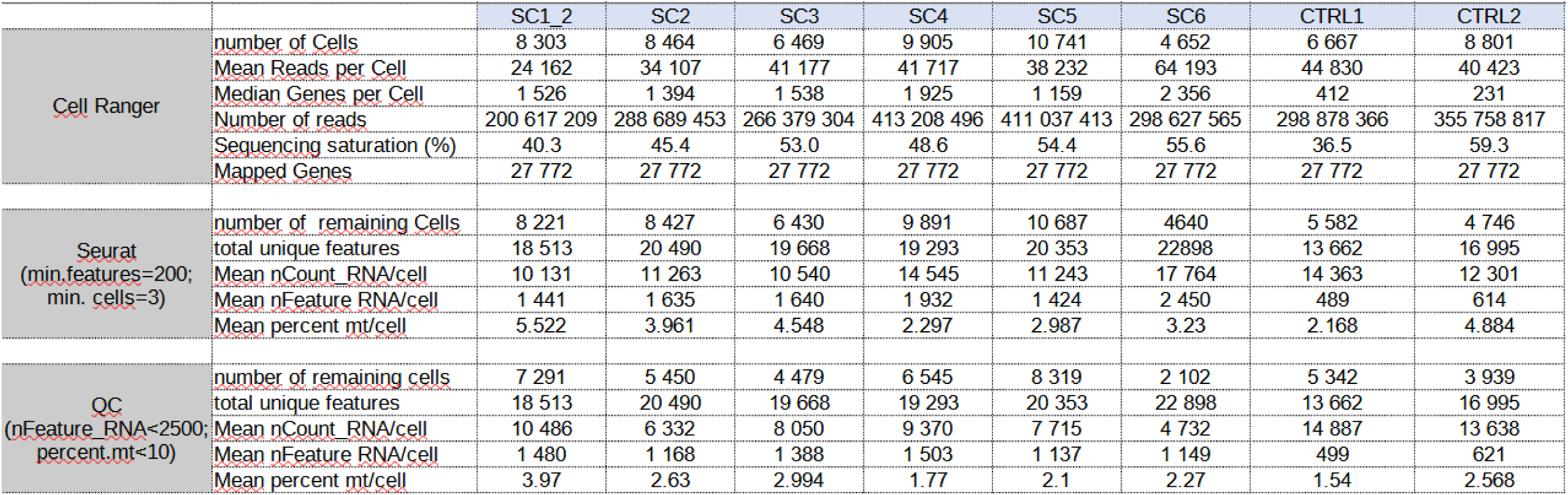
Table showing quality control metrics for single-cell RNA-seq data across all zebrafish samples (SC1–SC6, CTRL1–CTRL2).

**Supplementary Figure 6 :**
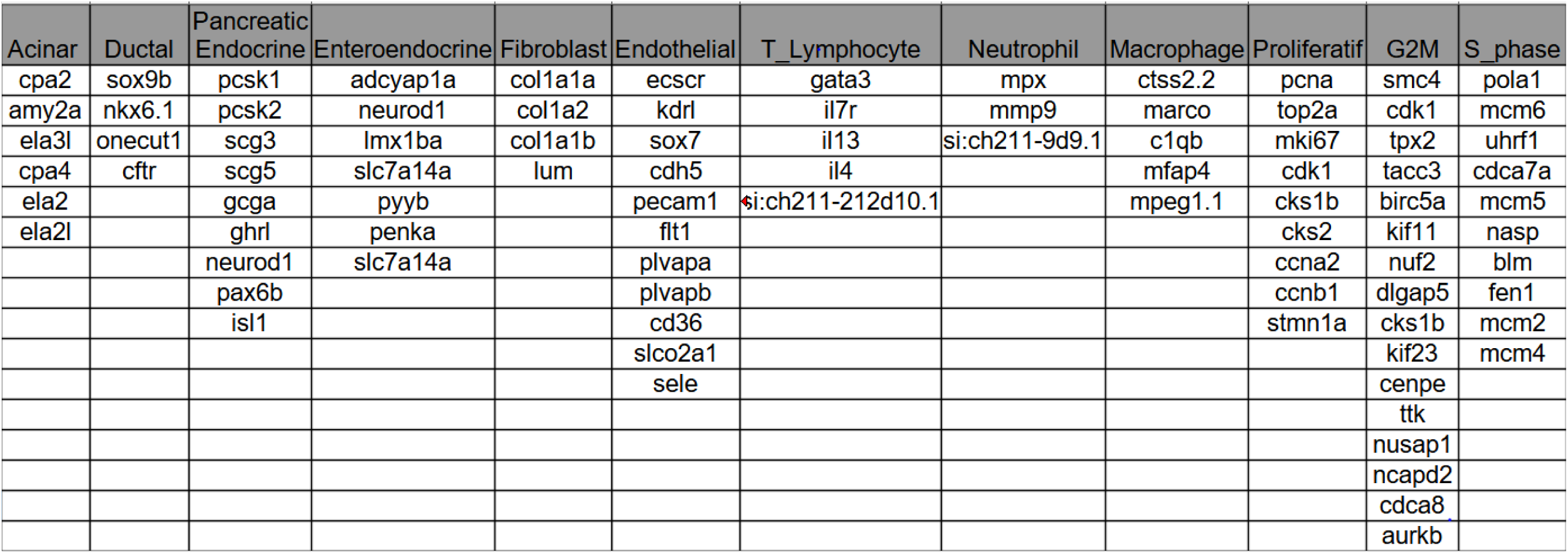
List of markers identified from literature and single-cell transcriptomic atlases of zebrafish development.

**Supplementary Figure 7 :**
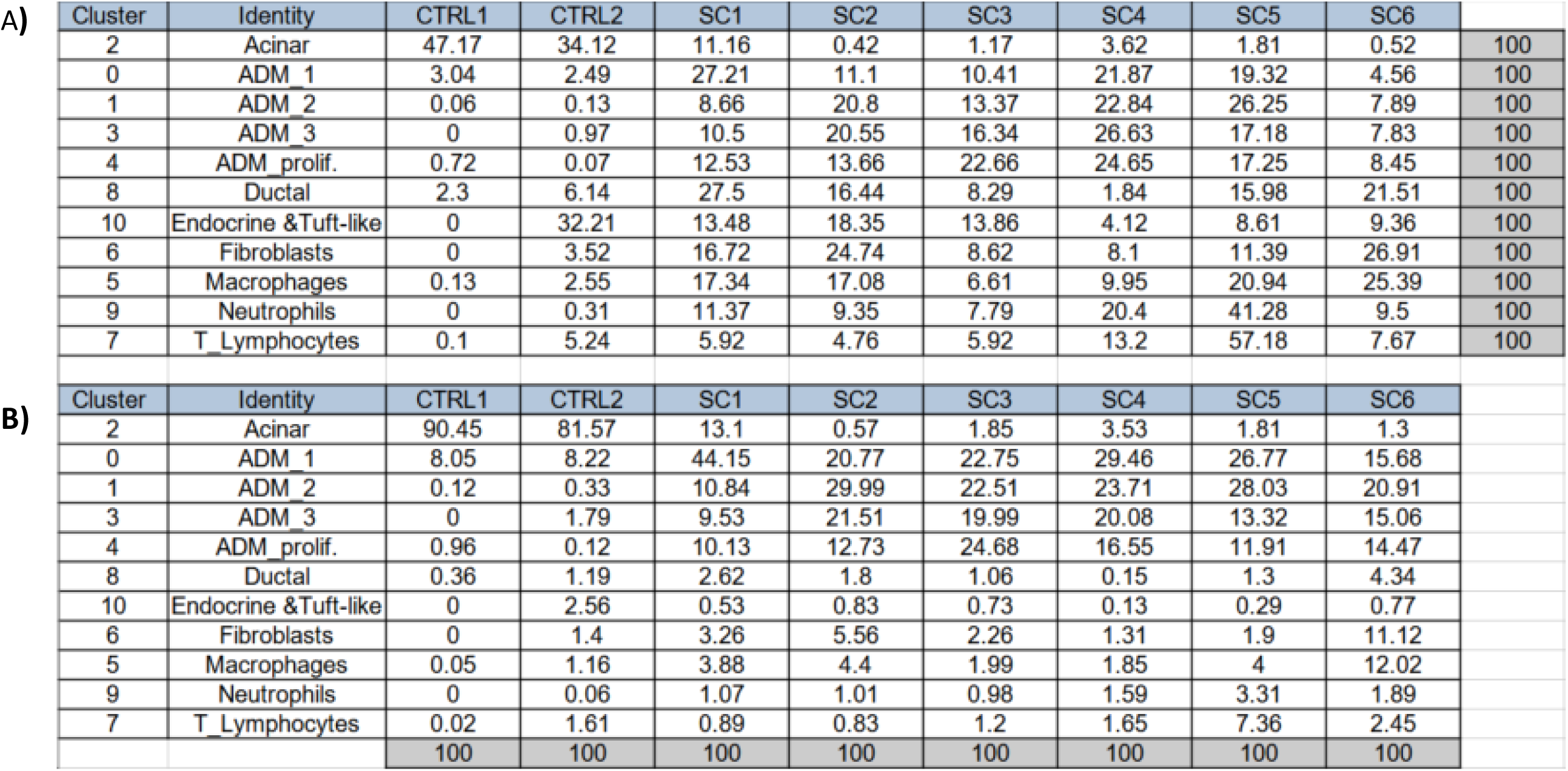
Sample distribution across cell clusters. Proportion of cells in each cluster derived from each individual sample. Sample distribution across cell clusters (A). Proportion of cells from each sample assigned to the indicated clusters **(B).**

**Supplementary Figure 8 :**
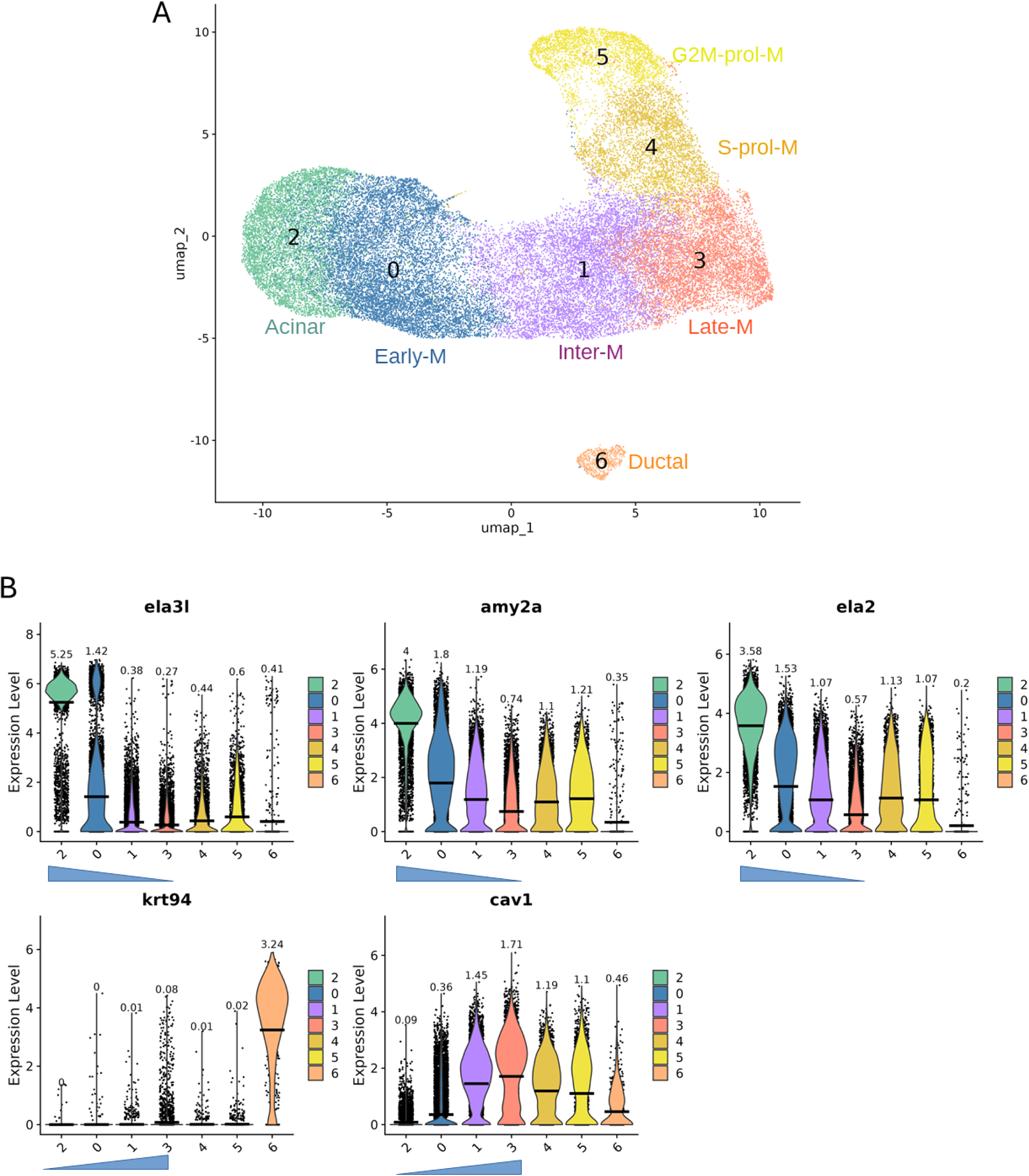
Gradual Loss of Acinar Identity and Induction of Metaplastic Markers in zebrafish metaplastic cells. A) UMAP representation of the zebrafish scRNAseq data restricted to pancreatic exocrine cells B) Violin plots illustrating the gradual loss of the acinar markers *ela3l*, *amy2al*, and *ela2*, accompanied by a progressive increase in the metaplastic markers *krt94* (orthologous to *KRT19*) and *cav1,* supporting a metaplastic trajectory from the acinar cl2 → cl 0 → cl 1 → cl 3. The mean expression level is shown as a horizontal bar, with the corresponding value indicated above each violin.

**Supplementary Figure 9:**
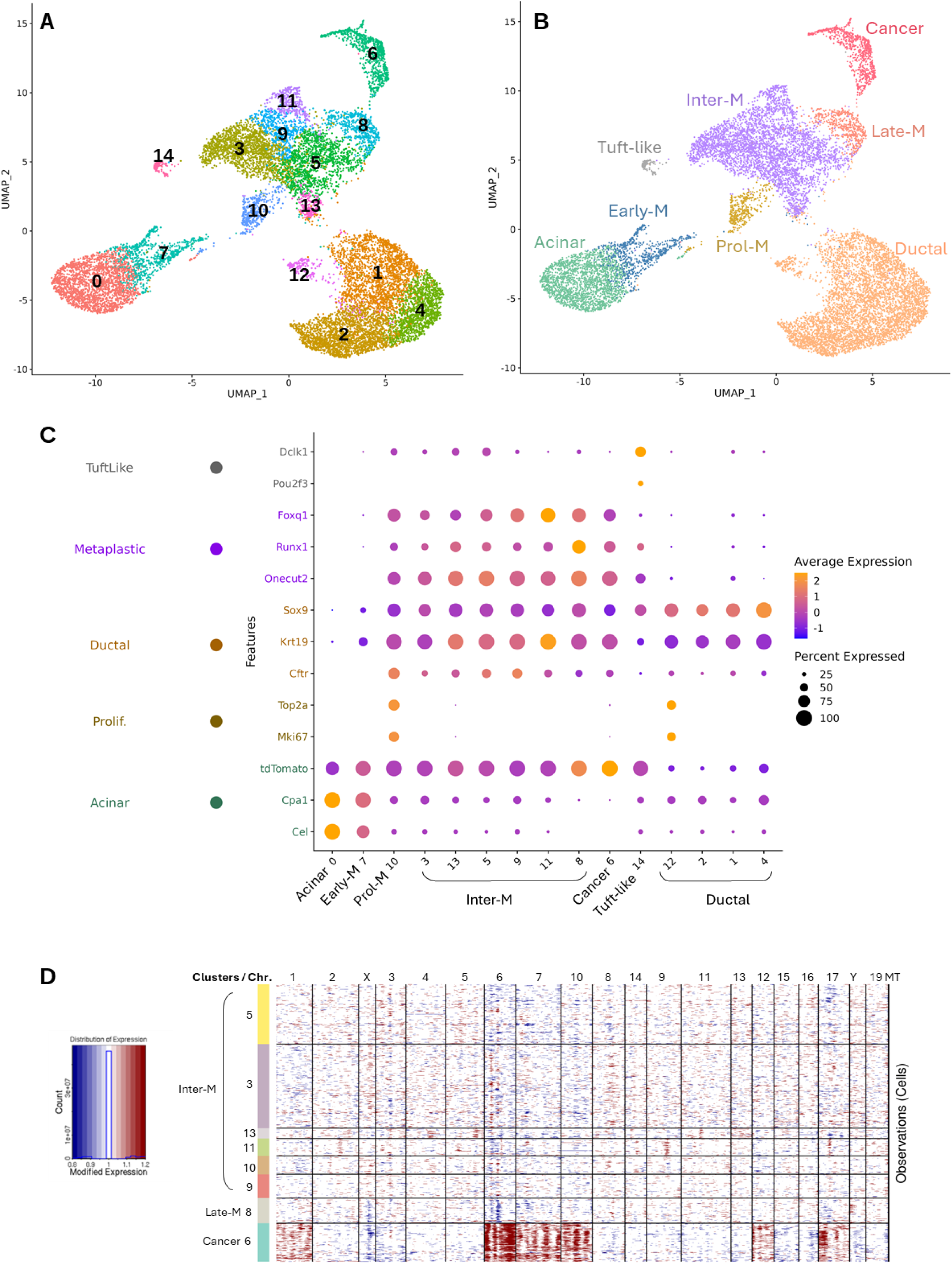
Characterization of murine scRNAseq data of pancreatic preneoplastic and cancer cells. scRNA-seq data from Schlesinger *et al.* were re-analysed [11]. Acinar-derived tumours were induced by expressing the oncogenic *Kras*^G12D^ variant in *Ptf1a-CreER; LSL-KRASG12D; LSL-tdTomato* mice, and pancreata were collected at six time points after tamoxifen induction (17 days, 6 weeks, 3, 5, 9, and 15 months). Only pancreatic cells were analysed, excluding the cells of the microenvironment**. A-B** Representation showing the distinct clusters of pancreatic cells (A), and their grouping according to their characteristics (B). **C:** Dot plot displaying the expression of cell type–specific markers across clusters. Dot size represents the proportion of cells expressing each gene, and color intensity reflects the average expression level among expressing cells. **D.** CNV analyses showing that the cancer cluster 6 exhibits extensive copy-number aberrations, whereas very few are detected in the late metaplastic cluster 8 and none in the other metaplastic clusters. On the x-axis, mouse chromosome regions and on the Y-axis, clusters numbers. The copy number variation score is indicated on the left.

**Supplementary Figure 10:**
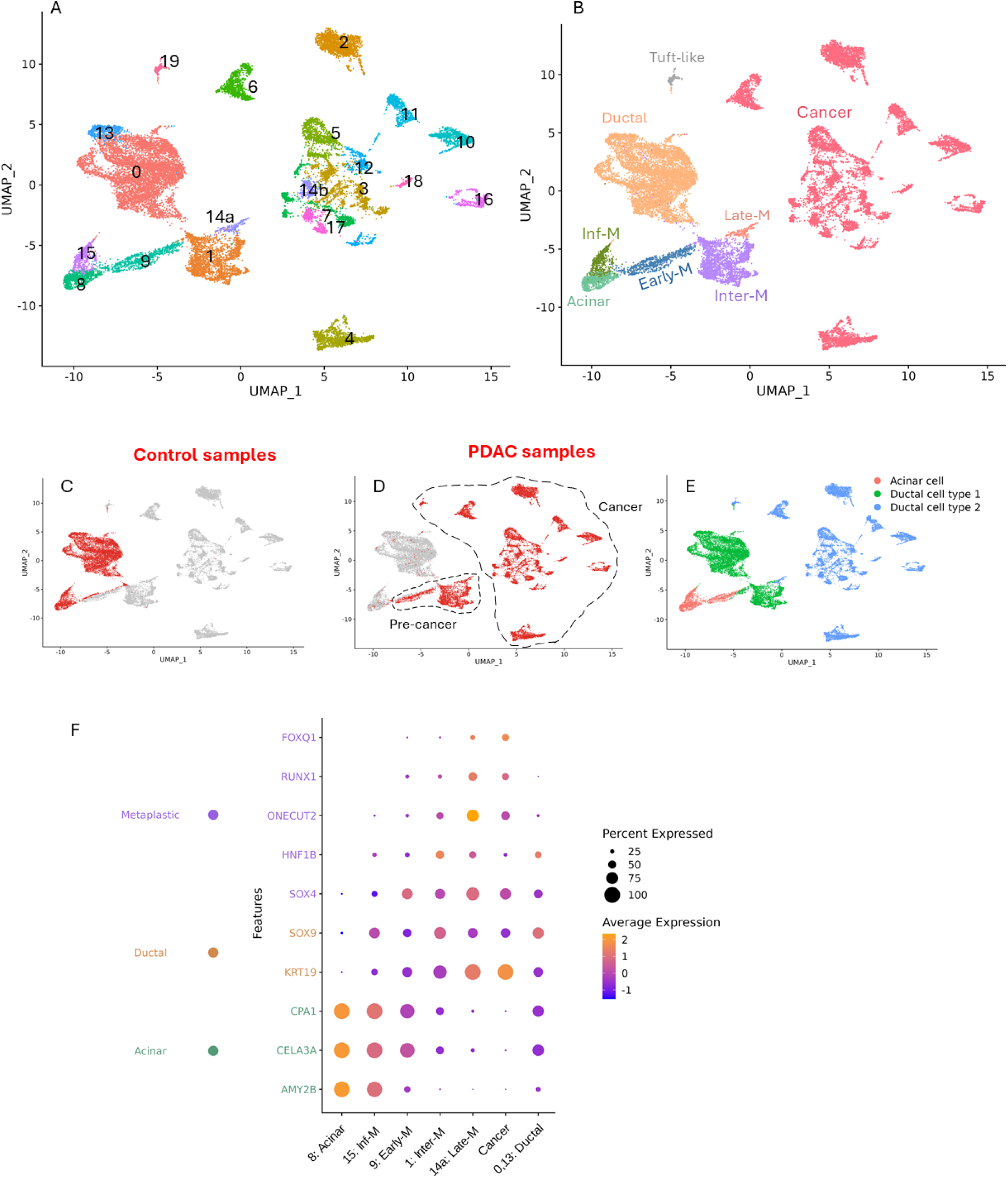
Characterization of scRNAseq data of human PDAC. Analyses of scRNA-seq data performed by Peng et al on PDAC samples from 24 patients and on pancreatic tissue from 11 control individuals suffering from non-pancreatic diseases [39]. Only pancreatic cells were analysed, excluding the cells of the microenvironment. **A–B.** UMAP representations showing the distinct clusters of pancreatic cells (A) and their grouping according to their characteristics (B). **C–D.** UMAP representation highlighting in red the contribution of control pancreatic cells (C) and patient-derived cells (D) relative to all other cells shown in grey. **E.** UMAP representation showing the localisation of acinar cells, ductal cell type 1 and ductal cell type 2 as identified by Peng et al. **F.** Dot plot displaying the expression of specific markers across clusters. Dot size represents the proportion of cells expressing each gene, and colour intensity reflects the average expression level among expressing cells.

**Additional File 1. Differentially expressed genes in metaplastic versus healthy acinar cells.** This table lists differentially expressed genes identified by comparing intermediate and late metaplastic cells to healthy acinar cells (absolute log2FC > 0.25, min.pct > 0.1, FDR < 0.001) in zebrafish (sheet “dr_DEG”), mouse (sheet “mm_DEG”), and human (sheet “hs_DEG”). The sheet “common_DEG_up” contains the list of genes upregulated in all three species. *pct1* refers to intermediate and late metaplastic cells, and *pct2* to healthy acinar cells. When genes were identified in all three datasets, the last column indicates the corresponding human ortholog.

**Additional file 2. Differentially expressed genes in the pancreatic multipotent progenitors.** This table lists the differentially expressed genes identified by comparing the transcriptome of ptf1a:GFP positive cells versus ptf1a:GFP negative cells. The sheet “DEG_up_GFP+” and DEG_down_GFP+ contains the list of genes upregulated and downregulated respectively in the pancreatic multipotent progenitors.

